# A rational design strategy and validation for protease-resistant fusion-inhibitor antiviral peptides

**DOI:** 10.64898/2026.09.12.750390

**Authors:** Kailu Yang, Chuchu Wang, Francesco Topi, Serena Muratcioglu, Ravi Kant, Tomas Strandin, Yih Tyng Bong, Lauri Kareinen, Sanna Maki, Saber H. Saber, Tobias Binder, Tarja Sironen, Subu Subramanian, K. Ian White, Richard A. Pfuetzner, Luis Esquivies, Olli Vapalahti, Merja Joensuu, John Kuriyan, Jussi Hepojoki, Giuseppe Balistreri, Axel T. Brunger

## Abstract

Peptide-based fusion inhibitors are promising pharmaceuticals in the fight against enveloped viruses relying on membrane fusion for host infection. However, peptide therapeutic applications have long been hindered by their poor stability *in vivo*. Here, we discovered that peptide inhibitors with the wildtype sequence of the heptad repeat 2 (HR2) domain of the SARS-CoV-2 spike protein are efficiently cleaved by Transmembrane Protease, Serine 2 (TMPRSS2), a key protease involved in the SARS-CoV-2 virus-cell fusion pathway. We then identified the corresponding cleavage sites and designed three protease-resistant peptides using ranking based on deep mutational scanning and natural occurrence. The three candidates all exhibit inhibitory activity in a cell-cell fusion assay. A high-resolution cryo-EM structure of the top candidate, HR2-NHN, bound to its HR1 target reveals the molecular basis for its potent activity. The top candidate of the cell-based screening assay showed significantly improved efficacy relative to the wildtype peptide when administered at different times before infection in both an authentic virus-cell infection assay and a mouse assay. More broadly, our results suggest that the design strategies for protease-resistant peptides could be applied to a broad spectrum of other enveloped viruses and pave the way for the development of safe, prophylactic antivirals that can be administered before exposure.

## Introduction

In enveloped viruses such as severe acute respiratory syndrome coronavirus 2 (SARS-CoV-2), membrane fusion is necessary for viral entry into the host cell^1,2^. Importantly, a conformational change within the spike protein (S) drives membrane fusion^3,4^, and peptide fragments of S derived from its heptad repeat 2 (HR2) motif bind its conserved heptad repeat 1 (HR1) motif^5^, effectively inhibiting fusion^6,7^. Indeed, HR2-based peptides have great potential as antivirals since they can be easily administered, for example, intranasally^8^ when an individual is expected to be at high risk of virus exposure. Furthermore, peptides without chemical modification are expected to exhibit minimal adverse effects, such as liver toxicity, compared with traditional small-molecule drugs^9^. However, a major concern regarding the therapeutic use of these peptides is poor *in vivo* stability. While chemical modifications such as lipidation^10,11^ and hydrocarbon stapling^12,13^ are among the common ways to minimize degradation, such modifications involve costly synthesis. Alternatively, improved peptide stability could be achieved by mutating protease cleavage sites likely to be encountered upon administration to the airways.

We sought to improve the stability of unmodified inhibitory HR2 peptides for SARS-CoV-2 *in vivo*. We discovered that HR2-based inhibitor peptides are efficiently cleaved by Transmembrane Protease, Serine 2 (TMPRSS2), a key protease that cleaves S at the S2′ site, allowing S to transition to the post-fusion state^6,14^. Based on this discovery, we hypothesized that the *in vivo* stability of HR2 peptides could be improved by mutating TMPRSS2 cleavage sites, since the peptides would likely encounter TMPRSS2-expressing host cells upon administration. After identifying the cleavage sites by mass spectrometry, we designed three triple mutant HR2 peptides resistant not only to TMPRSS2 cleavage but also to cleavage by the entire trypsin-like protease family, using rankings from both deep mutational scanning (DMS) and natural occurrence. All three protease-resistant design candidates exhibit inhibitory activity in a cell-cell fusion assay. We determined the high-resolution cryo-EM structure of the top candidate with the highest inhibitory activity (referred to as HR2-NHN) bound to HR1; this structure explains the effect of a key mutation, R1185H, on binding. Finally, tests in an authentic SARS-CoV-2 cell-infection assay and a mouse assay both validate HR2-NHN as an equally potent and more stable inhibitor compared to previously studied wildtype HR2 peptides.

Our study thus offers compelling new opportunities for prophylaxis or for ameliorating symptoms if administered soon after infection with SARS-CoV-2, which continues to evolve and spread, with four major variants (XFG, BA.3.2, PQ.16.1.1, and NB.1.8.1) currently under monitoring by the WHO. Variants of SARS-CoV-2 continue to challenge presently available therapies and pose a substantial health threat, including Long COVID^15^. Although these variants appear to have limited immune escape from the latest mRNA booster vaccines, an urgent need remains for orthogonal antiviral drugs that target relatively conserved regions of the virus. Such a strategy would lead to drugs that would remain effective against new variants, avoid large-scale immunization, be easily administered to prevent infection in advance or alleviate symptoms after infection, and have minimal side effects. More broadly, given widespread expression of trypsin-like proteases at mucosal and epithelial barriers^16–18^ and universal presence of basic residues within HR2-equivalent regions of many type I enveloped viruses (**Fig. S1**)^19–33^, our results inspire re-design of protease-resistant versions against a broad spectrum of other deadly viruses including Ebola virus^20^, Nipah virus^25^, respiratory syncytial virus (RSV)^21^, and human immunodeficiency virus (HIV)^19^.

## Results

### Widely used wildtype HR2 peptides are highly susceptible to TMPRSS2 cleavage, which abolishes their inhibitory activity

HR2 peptides inhibit SARS-CoV-2 infection, presumably by binding to the pre-hairpin intermediate of S following cleavage of S at the S1/S2 and the S2′ sites mainly by furin and TMPRSS2^6,34–36^. Although TMPRSS2 cleavage of the HR2 domain of the full-length S protein is not reported in any of its conformational states, we speculated that TMPRSS2 might cleave individual HR2 inhibitor peptides, considering that they are monomeric^6^, more accessible, and less structured than the HR2 domain within the context of full-length S. Therefore, we assessed the cleavage of three previously studied (**Fig. 1a**)^6,37^ fluorescently labeled HR2 peptides (42G_FL, longHR2_FL, shortHR2_FL) by TMPRSS2 using clear-native polyacrylamide gel electrophoresis (CN-PAGE) (**Fig. 1b**). All HR2 peptides tested were completely cleaved by TMPRSS2 after 1 h incubation at 37 °C, and cleavage was blocked by the serine protease inhibitor phenylmethylsulfonyl fluoride (PMSF) (**Fig. 1b**). Mass spectrometry revealed three fragments with two definitive cleavage sites at R1185 and K1191 and another possible cleavage site at K1181 (**Figs. S2 and S3**). We then asked if TMPRSS2-cleaved HR2 peptides could still inhibit SARS-CoV-2 fusion, using a previously established cell-cell fusion assay^6,38^. In this assay, ACE2 and S-expressing HEK cells were mixed, and content mixing was assessed with α-complementation of *E. coli* β-galactosidase. At variance to our previous studies, we developed a modified protocol to prepare solubilized peptides from the lyophilized state to achieve higher purity and more reliable results (see **Materials and Methods**). We found that after TMPRSS2 treatment, longHR2_42 loses inhibitory activity against the fusion mediated by the S protein of SARS-CoV-2 (**Fig. 1c**).

**Fig. 1.**
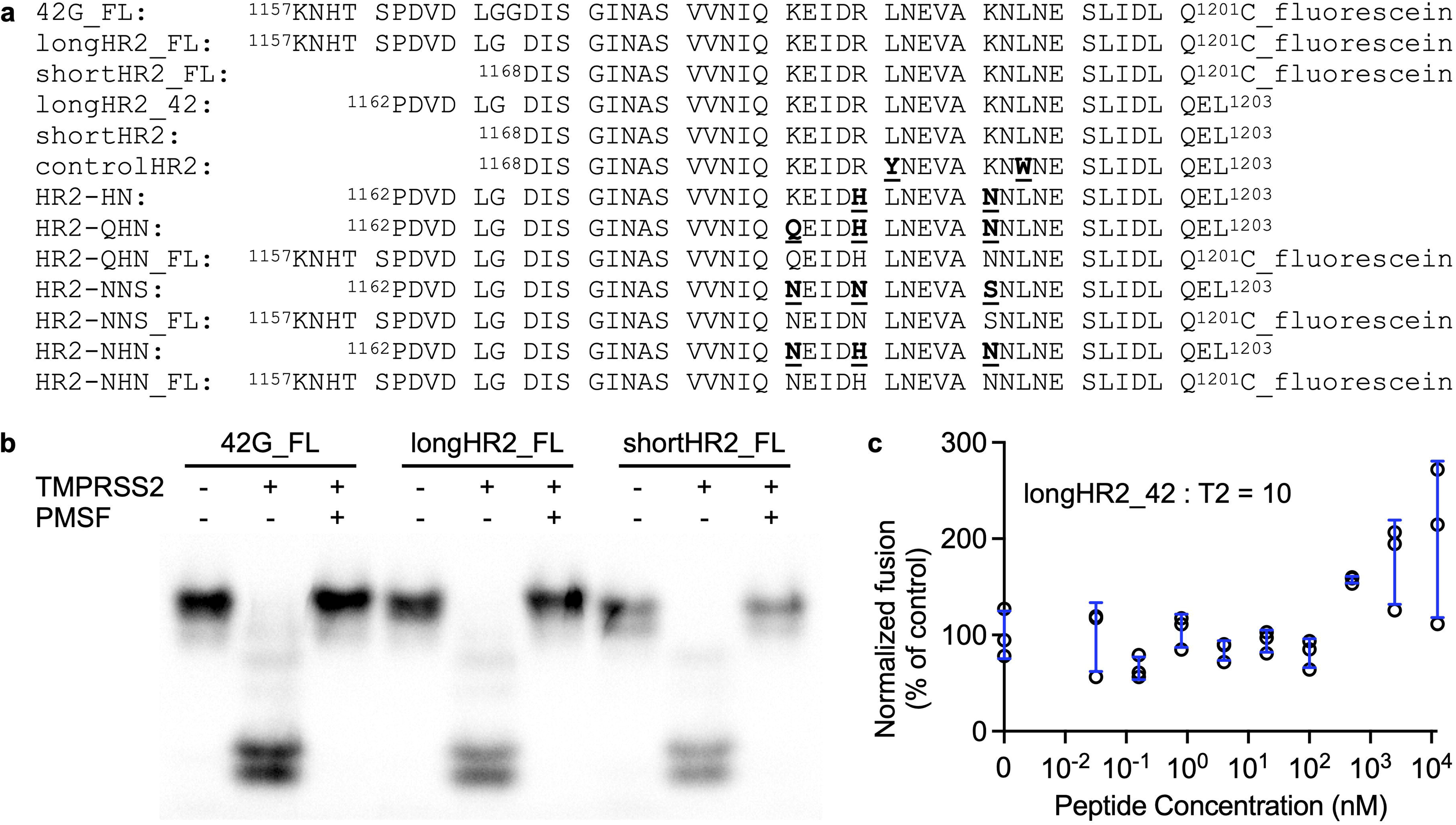
TMPRSS2 cleaves HR2 peptides and abolishes their inhibitory activity. (**a**) Primary amino acid sequences of HR2 peptides reported previously^6,37^ or in this study. The suffix “FL” indicates peptide labeling with fluorescein. For all other peptides without labeling, any difference from the wildtype sequence is indicated by an underline and bold font. (**b**) TMPRSS2 cleaves HR2 peptides as assessed by CN-PAGE fluorescence imaging. (**c**) Following TMPRSS2 cleavage, longHR2_42 does not inhibit fusion mediated by S, as assessed by a cell-cell fusion assay. The molar ratio between peptide and TMPRSS2 (T2) is marked in the panel. The raw data points are plotted as black circles, while the error bars (SD) are plotted in blue.

### Design of protease-resistant mutants based on deep mutational scanning and natural occurrence

Given these proteolysis results, we sought to design protease-resistant HR2 peptides. However, increasing peptide stability by mutagenesis potentially risks loss of antiviral efficacy, and the search space of possible mutations at three sites is large (20^3^ = 8000 sequences, including wildtype). To address these challenges, we used a two-pronged strategy to design stabilizing mutations at all three TMPRSS2-cleavage sites of HR2. First, we performed deep mutational scanning of the basic residues at the cleavage sites (K1181, R1185, and K1191) to rank all possible HR2 mutants by their ability to bind HR1 *in vitro* (**Fig. 2, a–b**). In this approach, a GFP-fused HR1 peptide is used as a bait to fish out the stronger binders from a library of *E. coli* cells that expose a mutant HR2 library at their surface; this mutant library includes all possible single mutations in the three basic residues of the HR2 peptide. After fluorescence-activated cell sorting (FACS), the bacteria that display the strongest fluorescence—i.e., the strongest binding to HR1-GFP—are subjected to deep sequencing to rank the most highly enriched mutations. Second, we surveyed the SARS-CoV-2 spike variants database^39,40^ to rank naturally occurring mutations of the three basic residues in the context of the full SARS-CoV-2 S protein (K1181, R1185, and K1191) (**Fig. 2c**). Among the ranked lists of possible mutations to non-basic residues (**Fig. 2, b– c**) we excluded mutations to hydrophobic residues to avoid potential solubility issues, and mutations to cysteine to avoid internal disulfide bonds or more extensive oxidative oligomerization. The deep mutational scanning data (**Fig. 2b**) suggest K1181N, R1185N, and K1191S (referred to as HR2-NNS, **Fig. 1a**) as a design candidate. Note that the introduction of most mutations at these positions was beneficial for HR1 binding; the majority of the HR2 variants bound more tightly than the wildtype peptide. On the other hand, the natural occurrence (**Fig. 2c**) suggests K1181N, R1185H, and K1191N (referred to as HR2-NHN, **Fig. 1a**) as a design candidate. In addition, we selected K1181Q, R1185H, and K1191N (referred to as HR2-QHN; **Fig. 1a**) as potential candidates (excluding the naturally occurring basic arginine and hydrophobic isoleucine at the K1181 position).

**Fig. 2.**
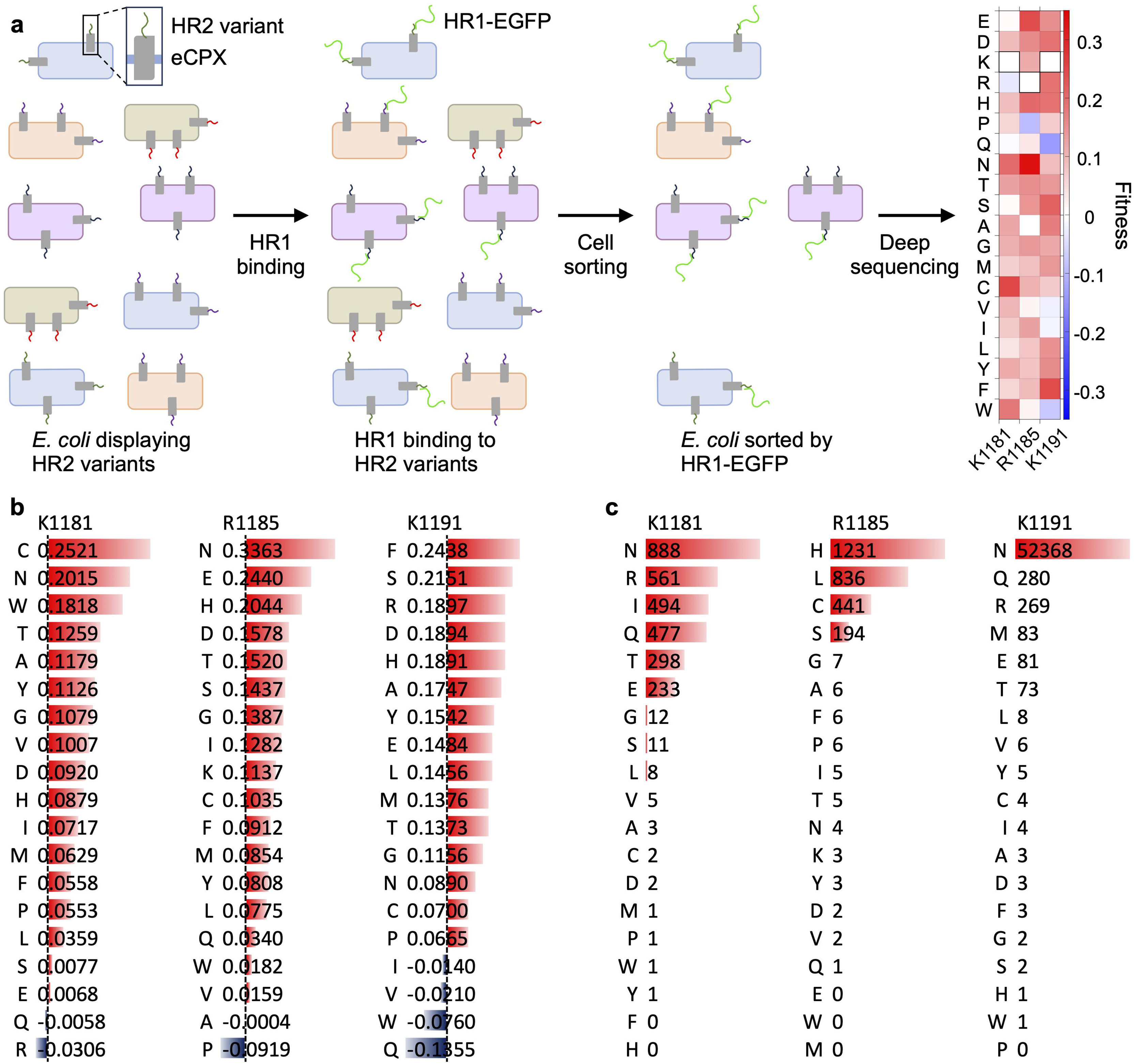
Ranking of mutations based on deep mutational scanning and natural occurrence. (**a**) Schematic representation of the high-throughput platform screening for HR1 binding. A library of *E. coli* cells displaying HR2 variants is mixed with GFP-labeled HR1 peptides to allow binding between HR1 and HR2. The cells are ranked based on GFP fluorescence and subjected to deep sequencing. Enrichment is calculated as the ratio of the number of sequencing reads for each variant in the sorted and unsorted populations (see **Materials and Methods** for details). All enrichment values are log_10_-transformed and normalized relative to the wildtype sequence, which has a value of 0. The effect of single amino acid substitutions of the three basic residues (K1181, R1185, and K1191) on HR1 binding is shown. Red and blue colors indicate that the specific mutant binds better or worse, respectively, than the wildtype sequence. (**b**) Enrichment rank for the three basic residues susceptible to TMPRSS2 cleavage of every single amino acid substitution ranked by relative enrichment, with the most beneficial mutations shown first. (**c**) All possible mutations for the three basic residues susceptible to TMPRSS2 cleavage ranked by natural occurrence. The number denotes the number of observations of each mutation in the GISAID-derived SARS-CoV-2 spike variants database^39,40^.

We prepared all three peptides (quality control data shown in **Figs. S4 to S6**) and verified that they are indeed resistant to TMPRSS2 cleavage (**Fig. 3a**).

**Fig. 3.**
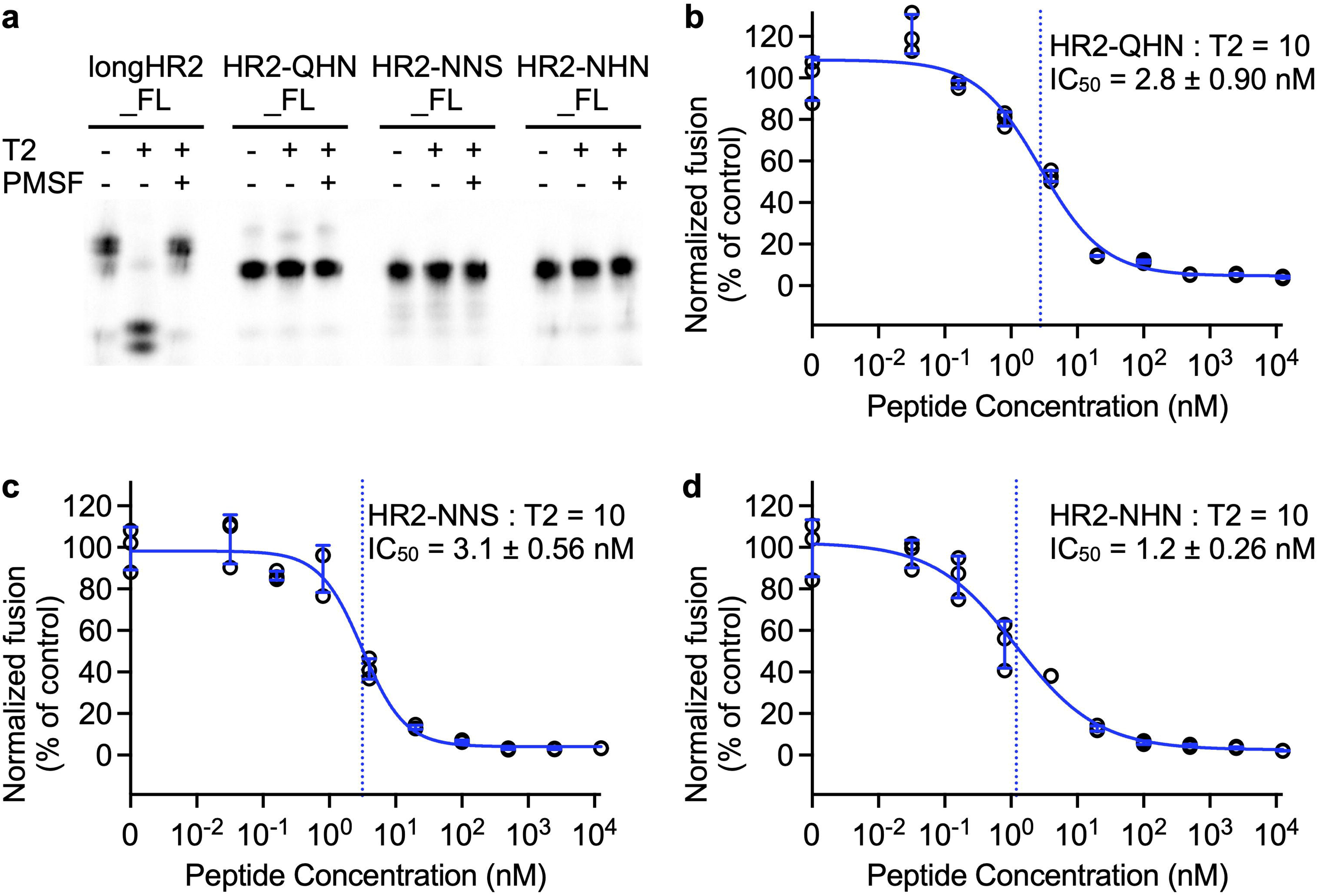
Cleavage assay and screening of the designed peptides. (**a**) The wildtype peptide (longHR2_FL) and the three newly designed mutant peptides (HR2-QHN_FL, HR2-NNS_FL, and HR2-NHN_FL, see Fig. 1a for their sequences) are labeled, subjected to TMPRSS2 (T2) cleavage, and then analyzed by CN-PAGE fluorescence imaging (see **Materials and Methods** for details). (**b–d**) Inhibitory activities of HR2-QHN (**b**), HR2-NNS (**c**), and HR2-NHN (**d**) in the cell-cell fusion assay. The molar ratio between peptide and TMPRSS2 (T2) is marked in the panel. The raw data points are plotted as black circles, while the error bars (SD), fitted curves, and vertical dashed lines at IC_50_ are plotted in blue.

### Screening of HR2 mutants by a cell-cell fusion assay

To assess the efficacy of the three designed peptides, we first tested their capacity to inhibit cell-cell fusion in the cell-cell fusion assay^6,38^ in the presence of TMPRSS2. We found the HR2-NHN peptide to have the highest inhibitory activity, with the other two peptides close behind (**Fig. 3, b–d**).

### The R1185H mutation disrupts a salt bridge but drives formation of an additional hydrogen bond between HR1 and HR2

Given the finding that the designed HR2 peptides bind HR1 so tightly, we sought to determine the molecular basis of these interactions, particularly because the basic R1185 of HR2 was previously shown to form a salt bridge with the acidic D936 of HR1. To do so, we performed single-particle cryo-EM of HR1 bound to the best-performing HR2-NHN using a previously reported molecular scaffolding method^41^ (**Fig. S7, Table S1**). The scaffolding method links the trimeric N termini of four HR1 fragments to four trimeric C termini of the Dps4 dodecamer from *Nostoc punctiforme*, and then complexes with HR2 peptides, enabling structure determination of the small HR1/HR2 complex by single-particle cryo-EM. While the R1185H mutation disrupts the salt bridge between R1185 of HR2 and D936 of HR1, the histidine side chain forms a compensatory hydrogen bond with S940 of HR1 (**Fig. 4**). In contrast, K1181 and K1191 do not show well-ordered sidechain density, consistent with the fact that they are surface-exposed and do not interact with HR1. After mutating these two lysine residues to asparagine, no new interaction with HR1 is formed, although the sidechain densities of the two asparagine residues are well-defined.

**Fig. 4.**
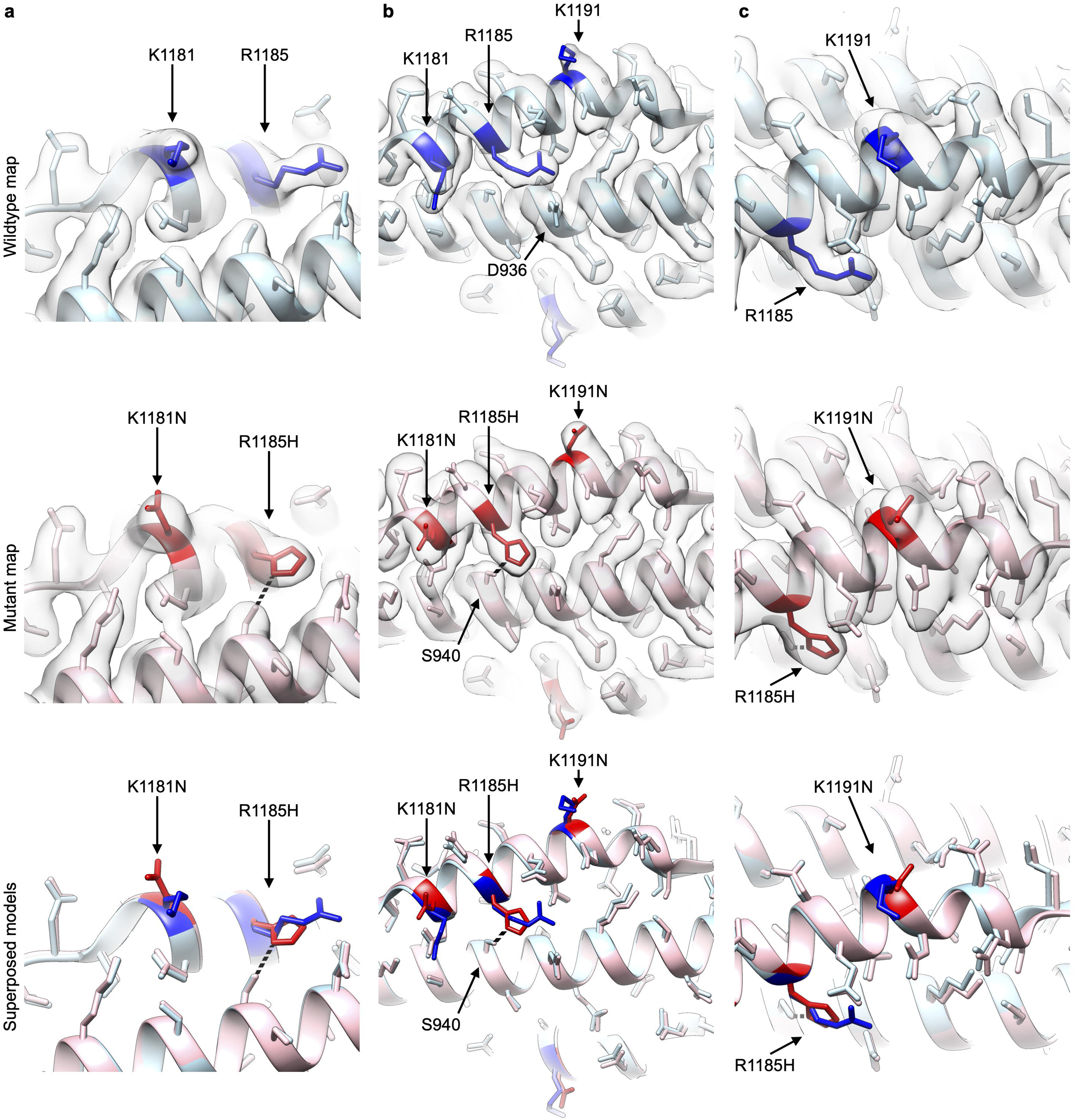
Structural insights into how the three mutations affect interaction with HR1. Comparison of cryo-EM structures of the wildtype longHR2_42 and HR2-NHN bound to HR1 in three different views. Panel (**a**) is focused on K1181N, panel (**b**) is focused on the entire helix of HR2, and panel (**c**) is focused on K1191N, with R1185H visible nearby in all three panels. The wildtype cryo-EM structure (light blue) and map are shown in the top row, the HR2-NHN (pink) structure and map are displayed in the middle row, and the superposed wildtype and HR2-NHN models without maps are displayed in the bottom row. The mutated residues are colored blue for the wildtype structure and red for the mutant structures. The hydrogen bond between H1185 and S940 is shown as a black dashed line in the middle and bottom rows for panels (**a–c**).

### The TMPRSS2-resistant HR2-NHN maintains inhibitory activity longer than the wildtype longHR2_42 in an authentic SARS-CoV-2–cell infection assay

Protease-resistant peptides are expected to have a longer bioavailability when proteases such as TMPRSS2 or cathepsins are present at the cell surface or in the endosomal compartment of cells *in vitro* or *in vivo*. To test if the antiviral activity of the best-performing TMPRSS2-resistant peptide HR2-NHN persisted longer than the original peptide longHR2_42, we preincubated Calu-3 cells, a lung-derived human line endogenously expressing ACE2 and TMPRSS2^42^, with a serial dilution of the respective peptides for 72 h or 0 h (*i.e.*, just before infection) followed by infection for 18 h with a recombinant SARS-CoV-2 Wuhan-Hu-1 (Wuh) strain expressing the mCherry red fluorescent protein^43^ for detection by high-content fluorescence microscopy (**Fig. 5**). When administered just before infection (0 h), both peptides inhibited infection in a dose dependent manner and with similar IC_50_s (**Fig. 5, a–b**). Preincubation with the cells for 72 h prior to infection resulted in a decrease in the antiviral activity of the original longHR2_42 peptide, the IC_50_ of which shifted from 2.6 nM to 8.9 nM, but not for the protease-resistant HR2-NHN, which retained its antiviral activity (IC_50_ 1.9 nM) (**Fig. 5, c–d**). Statistical analysis of the area under the curve (AUC) confirmed that this difference was significant (**Fig. S8, a–b**). Both peptides also inhibited the Omicron variant XBB.1.5^44^, but with lower potency than the Wuh strain (**Fig. 5, e– f, and Fig. S8c**). The decreased potency against Omicron was expected, as we have previously shown that mutations in the HR1 region of the Omicron spike protein impair binding of longHR2_42^37^.

**Fig. 5.**
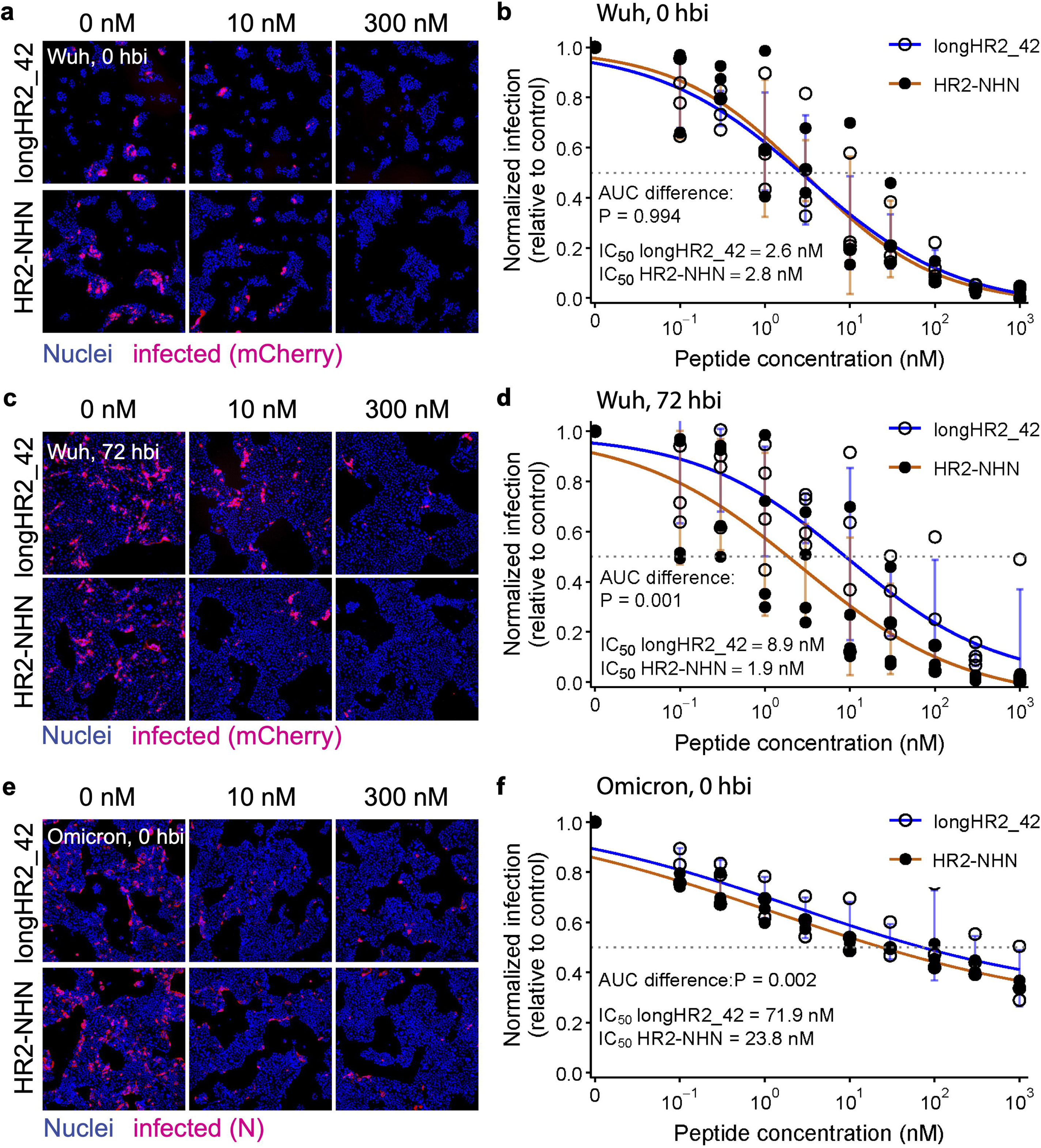
The TMPRSS2-resistant HR2-NHN maintains inhibitory activity longer than wildtype longHR2_42 in an authentic SARS-CoV-2 cell infection assay. Representative fluorescence images (**a, c, e**) and quantification after automated image analysis (**b, d, f**) of Calu-3 cells treated with indicated doses of longHR2_42 or HR2-NHN at 0 h (**a–b and e– f**) or 72 h (**c–d**) before infection (hbi) for 18 h with the SARS-CoV-2-mCherry Wuhan-Hu-1 strain (**a–d**) or Omicron XBB.1.5 (**e–f**). The estimated IC_50_ of the two peptides after different preincubation times and the statistical significance (*P*) of the difference between the Area Under the Curve (AUC) for each peptide are indicated in the graphs (see also **Fig. S8**). Individual points represent independent repetitions; error bars indicate SD. Solid lines show the fitted dose-response curves. The horizontal dotted line denotes normalized infection of 0.5. Blue curves, longHR2_42; orange curves, HR2-NHN. IC_50_, curve fitting, and respective AUCs were determined using R. Statistical analysis was also performed in R using the bootstrap resampling method for dose and time responses (see **Materials and Methods**).

Supporting the results obtained in Calu-3 cells, longer bioavailability for HR2-NHN compared to longHR2_42 was also observed in A549 cells stably over-expressing human ACE2 and TMPRSS2 (named A549-AT)^45^ (**Fig. S8**). In these experiments, longHR2_42 and HR2-NHN were each tested at a fixed concentration of 300 nM and added 0, 12, or 24 h before infection.

Over 50% of the inhibitory activity of the protease-resistant peptide HR2-NHN against SARS-CoV-2 Wuh was retained following preincubation for 12 h, while, in marked contrast, the inhibitory activity of the wildtype longHR2_42 is completely lost for the same period of preincubation (**Fig. S8, d–e**). With a preincubation period of 24 h, both peptides lose all inhibitory activity, probably because the peptides are either endocytosed into the cells after 24 h or degraded by multiple host proteases in A549-AT cells.

### The TMPRSS2-resistant HR2-NHN maintains antiviral activity longer than the wildtype longHR2_42 in SARS-CoV-2-infected mice

To test if HR2-NHN also had improved bioavailability *in vivo* compared to longHR2_42, we measured their activity in wildtype BALB/c female mice with the Beta variant of SARS-CoV-2, which infects wildtype mice and causes lung infection within 24 h^46^.

First, to determine the optimal peptide concentration to use *in vivo* before a direct comparison of longHR2_42 and HR2-NHN, we tested in mice (4 mice per treatment group) two concentrations of longHR2_42, 1 mg/kg and 0.1 mg/kg and administered the peptide just before infection (0 hbi, 1 mg/kg and 0.1 mg/kg), 3 h before infection (3 hbi, 1 mg/kg), or 3 h post infection (3 hpi, 1 mg/kg) (**Fig. 6a**). The peptides (or PBS control) and the virus (2 × 10^5^ PFU/mouse) were administered intranasally as 20 μL intranasal inoculations. In this first test, a second inoculation of the peptide was performed at 24 h post infection (hpi) (**Fig. 6a**), and after anesthesia at 48 hpi, mice were culled and RNA extracted for qRT-PCR analysis (**Fig. 6a**, end).

**Fig. 6.**
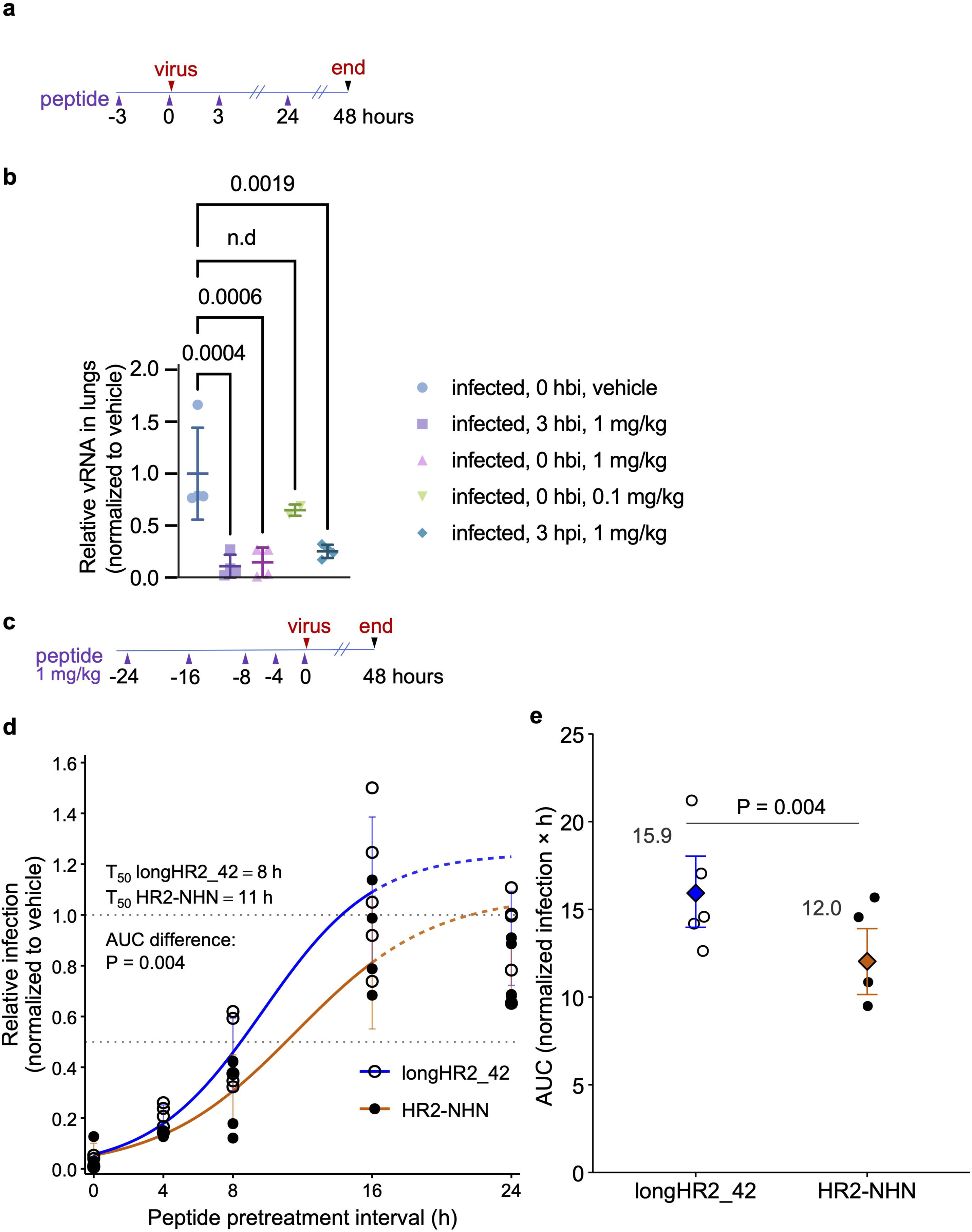
HR2-NHN peptide maintains antiviral activity longer than the wildtype longHR2_42 in SARS-CoV-2 infected mice. (**a**) Workflow of *in vivo* experiment identifying a protective dosing regimen for longHR2_42. (**b**) Relative lung viral RNA (vRNA), normalized to the mean of the vehicle-treated group (defined as 1.0), for mice treated with vehicle or longHR2_42 at the indicated pretreatment or post-infection time and doses. Individual points represent individual mice; the group mean and SD are indicated for each treatment; hbi, hours before infection; hpi, hours post infection. The P values indicating the statistical significance of each sample relative to vehicle-control treated mice are indicated in the graph. Statistical analysis was performed with Ordinary one-way Anova Dunnett’s multiple comparison test. For the mouse treatment “0 h 0.1 mg/kg”, two of the four mice died during anesthesia prior to infection; thus, statistical analysis was not performed for these mice (n.d.). (**c**) Workflow of an *in vivo* experiment to determine the duration of antiviral activity of the peptides with a dose of 1 mg/kg. (**d**) Re-infection assay with viruses extracted from lungs of mice pretreated with peptides as described in **c** and used to infect VeroE6 cells for 18 h, followed by immunofluorescence and image analysis to identify infected cells. Individual points represent infections with viruses extracted from individual mice (see **Materials and Methods** for details). Error bars indicate SD. Infection was normalized to the vehicle-treated controls (defined as 1.0). Solid lines show the fitted time-response curves over 0–16 h; dashed lines indicate extrapolation beyond the fitted interval to 24 h. Horizontal dotted lines denote normalized infection values of 1.0 and 0.5. Estimated T_50_ values and the exact P value for the 0–24 h AUC comparison are indicated. n = 5 mice per peptide at each time point. Blue, longHR2_42; orange, HR2-NHN. **(e)** Area under the curve. AUC was calculated from the observed mean normalized infection values from each time point in (**d**). Diamonds indicate the group AUC estimates and error bars indicate bootstrap 95% confidence intervals. The exact bootstrap P value for the comparison between longHR2_42 and HR2-NHN is shown. Blue, longHR2_42; orange, HR2-NHN.

When the longHR2_42 was administered intranasally at 1 mg/kg at 0 hbi (*i.e.*, a few seconds before intranasal virus inoculation), infection in the lungs was strongly inhibited as determined by qRT-PCR of RNA extracted from the lungs at 48 hpi (**Fig. 6b**, 0 hbi, 1 mg/kg). This inhibition was dose-dependent, and at a dose of 0.1 mg/kg the inhibitory effect was no longer significant (**Fig. 6b**, 0 hbi, 0.1 mg/kg). Significant inhibitory activity was observed at a concentration of 1 mg/kg when the longHR2_42 peptide was administered prophylactically 3 hbi (**Fig. 6b**, 3 hbi, 1 mg/kg). To assess whether longHR2_42 would have antiviral activity when administered after virus infection (i.e., therapeutic activity), the peptide was administered at 3 h post-virus inoculation. Although to a lesser extent compared to prophylactic administration, significant antiviral activity was still observed (**Fig. 6b**, 3 hpi).

As a first comparison between the original and the TMPRSS2-resistant peptide, mice were pretreated at 0, 12, and 24 h before infection at a concentration of 1 mg/kg, as a single dose at the indicated times. The second dose at 24 hpi was not administered. The extent of lung infection was determined by qRT-PCR at 48 hpi as described above (**Fig. S9a**). Immunohistochemistry of mouse lungs from each treatment group was used to confirm the PCR results (**Fig. S9b**). Confirming the *in vitro* results, the antiviral activity of HR2-NHN was significantly higher than longHR2_42 when the peptides were administered 12 h before infection, as assessed by qRT-PCR (**Fig. S9c**, −12 hbi) and confirmed by a qualitative reduction in N protein staining (**Fig. S9b**, −12 hbi). However, when administered at a single dose 24 hbi, both peptides lost antiviral activity (**Fig. S9b**, −24 hbi).

To estimate the half-life of the two peptides’ antiviral activity, a more extensive time course (0, −4, −8, −16, −24 hbi) was performed in mice (**Fig. 6c**). While quantification of viral RNA by qRT-PCR is a sensitive method, it does not distinguish between intracellular viral genomes (or RNA transcripts) and extracellular infectious viruses, which are relevant to the spread of infection in the respiratory tract and to host-to-host transmission. To assess the extent of residual infectious viruses in the lungs of mice infected and treated with either vehicle control or peptides, at 48 hpi, mice were sacrificed, and lung homogenates were used to re-infect highly susceptible VeroE6 cells. As described above for the cell culture assays, at 18 hpi, infection rates were quantified by immunofluorescence with antibodies raised against the viral N protein, followed by automated fluorescence imaging and image analysis. Under these conditions, the apparent T_50_ of the original peptide longHR2_42 was approximately 8 h, while that of the protease-resistant HR2-NHN was 11 h (**Fig. 6d**). Statistical analysis of the AUC confirmed that these differences were statistically significant (**Fig. 6e**). If administered more than 16 h prior to infection, none of the peptides retained inhibitory effects (**Fig. 6d and Fig. S9c**).

## Discussion

Peptide therapeutics are notorious for their poor stability *in vivo*, and efforts to improve their stability and bioavailability are thus critical. Here, we describe a strategy to achieve this in the context of proof-of-concept peptide therapeutics targeting SARS-CoV-2. We observed that wildtype HR2-based inhibitors can be efficiently cleaved by TMPRSS2 (**Fig. 1b**), a key protease in the infection process. We therefore developed TMPRSS2-resistant peptides with vastly improved protease resistance and longer-lived potency through a strategy drawing on deep mutational scanning (**Fig. 2, a–b**) and natural occurrence (**Fig. 2c**), rationalized binding through structural studies with cryo-EM (**Fig. 4 and Fig. S7**) and tested our designs using multiple *in vitro* and *in vivo* approaches (**Figs. 3, 5, and 6**). The most efficacious protease-resistant peptide, HR2-NHN, is an HR2-based inhibitor with improved bioavailability that resists TMPRSS2 cleavage and outperforms its wildtype predecessor (longHR2_42) when administered at different times before infection in a mouse assay (**Figs. 5, 6 and Fig. S8**).

We used two ranking schemes, deep mutational scanning and surveying of natural occurrence (**Fig. 2**), for the design of three TMPRSS2-cleavage resistant peptides, HR2-QHN, HR2-NNS, and HR2-NHN (**Fig. 1a**). All three designs exhibit inhibitory activity (**Fig. 3**). The peptide HR2-NHN is the top candidate from the natural occurrence approach, and HR2-NNS is the top candidate from the deep mutational scanning approach. A difference between these two top designs is the choice of asparagine or histidine at position 1185, and the histidine mutant exhibits higher inhibitory activity. Combining different ranking schemes increases the exploration of possible mutants. In this context, we note the features of deep mutational scanning. First, it focuses on the binding between HR1 and HR2 and is not biased by other restraints that likely dominate some mutations in natural occurrence. For example, some frequently occurring mutations may be advantageous for prefusion conformations of the spike protein, or simply for improved viral genome stability, rather than for tighter binding between HR1 and HR2. Second, it theoretically works for any virus with a similar entry mechanism, even with limited sequencing data for a novel or rare virus.

Furthermore, although the peptides were designed to resist cleavage by TMPRSS2 specifically, we mutated all three basic residues. Therefore, the resultant HR2-NHN peptide is likely resistant to the entire trypsin-like protease family, which requires a basic residue at the P1 position for cleavage^47^. This could generally enhance bioavailability and help explain the contrast between HR2-NHN and wildtype longHR2_42 when preincubated with cells at different times before infection and when delivered intranasally in a mouse model of SARS-CoV-2 infection, even with a single, relatively low dose of 1 mg/kg. Interestingly, HR2-NHN also showed a modest advantage over longHR2_42 against Omicron XBB.1.5 without preincubation, as reflected in the overall dose-response AUC (**Fig. 5 and Fig. S8**). Omicron variants have been reported to favor endosomal entry over TMPRSS2-dependent cell-surface entry in several cellular contexts^48^. The strain-specific difference in entry route may influence the relative activity of the two peptides, particularly because the new peptide has all basic residues mutated, and the substantially altered overall net charge may lead to more efficient endocytosis together with the virus, as well as render it more resistant to endosomal degradation during the time of virus cell entry.

Our *in vivo* results indicate that intranasal delivery of monomeric, chemically unmodified fusion inhibitory peptides has both prophylactic and therapeutic potential. The described approach to confer protease resistance through rational mutagenesis extended the half-life of the respective peptide’s antiviral activity. This approach can be applied to fusion-inhibitor peptides raised against different viruses, such as other coronaviruses, influenza virus, and respiratory syncytial virus.

A recent study investigating the effects of PEGylation and lipidation on the bioavailability of SARS-CoV-2 inhibitor peptides estimated that unmodified peptides lose potency after approximately 8 h of administration, a finding that correlated with decreased plasma concentrations in treated mice^49^. Our study is consistent with this estimate and demonstrates that the mutagenesis strategy presented here can improve peptide stability *in vivo* without the need for chemical modifications, a significant advantage given the simplicity of manufacturing these monomeric peptides for potential therapeutic use.

We note that this is the first time that our unmodified peptides with N-terminal extension (the wildtype longHR2_42 and the mutant HR2-NHN) were studied in mice. When the peptides were administered simultaneously with the virus, or even three hours prior to infection, we observed complete blockage of infection with a single low dose at 1 mg/kg, compared to 4–20 mg/kg from a recent study using a dimer of shortHR2 with linked PEG and cholesterol^49,50^. Several factors may have contributed to the significant difference in potency. First, the N-terminal extension of the peptide likely plays a pivotal role in competing with the HR2 domain of S for binding to the HR1 domain. Second, although the linked PEG and cholesterol moieties offer several advantages, including increased solubility, improved stability, and enhanced membrane binding to increase local concentration, they also increase the overall mass of the inhibitor, potentially affecting efficacy. Third, we used wildtype mice and the Beta variant of the virus, and infection is milder than in mice transgenically expressing human ACE2 and thus easier to inhibit. Nevertheless, these mutations presented here could also be incorporated into other HR2-based designs.

Considering its superior stability, we envision that the new HR2-NHN inhibitor from this study could become a better prototype than the wildtype peptide for further tests and improvements to further enhance the efficacy of HR2-NHN. In particular, applying chemical modifications such as lipidation and hydrocarbon stapling might further improve the stability and potency of HR2-NHN^10–13^. Finally, we propose that our strategy could be readily performed for other enveloped viruses that usually contain basic residues in their HR2-equivalent regions (**Fig. S1**)^19–33^, including Ebola virus^20^, Nipah virus^25^, RSV^21^, and HIV^19^. Moreover, peptides, or protein binders, that interfere with entry factors of non-enveloped viruses could also be designed.

## Materials and Methods

### Purification of TMPRSS2

The purification of the secreted, C-terminally His-tagged TMPRSS2 ectodomain (residues 109– 492) was performed following a previously established protocol^36^. Briefly, the plasmid was obtained from Addgene (plasmid no. 176412). Expression was carried out in Sf9 insect cells using a baculovirus system. The culture supernatant, containing the secreted protein, was harvested 4–5 days post infection when cell viability reached 55–60%. After clarification by centrifugation, the supernatant pH was adjusted to 7.4, and the protein was captured via batch incubation with Ni-NTA resin. The resin was then collected, transferred to a gravity-flow column, washed with three column volumes of ice-cold PBS, and the protein was eluted using PBS containing 500 mM imidazole. For zymogen activation, the eluate was concentrated and dialyzed against an activation buffer (25 mM Tris pH 8.0, 75 mM NaCl, 2 mM CaCl_2_) for 6 h at room temperature. The final purification step involved size-exclusion chromatography on a Superdex 75 column equilibrated in 50 mM Tris pH 7.5, 250 mM NaCl. Fractions corresponding to the main protein peak were verified by SDS-PAGE with and without DTT, pooled, and concentrated with a 10 kDa MWCO concentrator (EMD Millipore cat. no. UFC901008). The SEC profile and SDS-PAGE (**Fig. S10**) were essentially the same as the protocol we followed^36^. For long-term storage, the purified active enzyme was prepared in a buffer containing 25% glycerol (50 mM Tris pH 7.5, 250 mM NaCl, 25% glycerol), flash-frozen in liquid nitrogen, and stored at −80 °C.

### Fluorescent labeling of peptides

The HR2 peptides for fluorescent labeling were prepared in-house as described before^6^. Briefly, HR2 peptides ranging from either residues 1157–1201 (longHR2_FL), or residues 1168–1201 (shortHR2_FL), or residues 1157–1201 with the glycine insertion (42G_FL)^37^, of the SARS-CoV-2 Wuhan-Hu-1 strain (Global Initiative on Sharing Avian Influenza Data [GISAID] accession ID: EPI_ISL_402124, here referred to as Wuh) S protein were cloned into the pETDuet-1 plasmid with N-terminal hexa-histidine and SUMO tags and a C-terminal cysteine for maleimide labeling. The peptides were recombinantly expressed in *E. coli* BL21(DE3) at 37 °C overnight using autoinducing LB medium^51^. Cells were harvested, resuspended in lysis buffer (50 mM Tris pH 8, 300 mM NaCl, 20 mM imidazole, 0.5 mM TCEP), and lysed by sonication. Lysate was clarified by centrifugation at 40,000 RPM for 30 min. Supernatant was bound to Nickel-NTA resin for 45 min at 4 °C with stirring. After washing with the lysis buffer, the bound protein was eluted with the lysis buffer supplemented with 300 mM imidazole. The eluted protein was then subjected to SUMO protease cleavage, heat treatment at 95 °C for 10 min, and SEC using a Superdex 30 HiLoad 16/60 column equilibrated with 20 mM sodium phosphate, pH 7.4, 150 mM NaCl, and 0.5 mM TCEP. The purified peptides were concentrated to ∼1 mM with a 2 kDa MWCO concentrator (Sartorius cat. no. VN02H91), flash frozen in liquid nitrogen, and stored in a −80 °C freezer for future use.

The maleimide labeling of the HR2 peptides was performed by mixing 100 μM peptide and 2.5 mM fluorescein-5-maleimide (Thermo Fisher, cat. no. 62245) in degassed buffer (50 mM sodium phosphate, pH 6.8, 150 mM NaCl, 0.05 mM TCEP), followed by overnight incubation at 4 °C. Following labeling, free dyes were removed by SEC using a Superdex 75 HR 10/300 GL column in PBS buffer (137 mM NaCl, 2.7 mM KCl, and 10 mM phosphate, pH 7.4). Labeled peptides were then further purified by high-performance liquid chromatography (HPLC) with a C18 column in a gradient of water and acetonitrile supplemented with 0.1% trifluoroacetic acid, followed by lyophilization and a final round of SEC using a Superdex 75 HR 10/300 GL column equilibrated with PBS buffer to remove any residual trifluoroacetic acid. The purified peptides were confirmed by mass spectrometry, concentrated to ∼50 μM, protected from light, and stored at −80 °C.

### TMPRSS2 cleavage of the fluorescently labeled peptides

Concentrations used for the cleavage were 1.5 μM for the fluorescently labeled peptides (42G_FL, longHR2_FL, shortHR2_FL), 0.3 μM for TMPRSS2, and 5 mM for PMSF, respectively. After mixing on ice, the cleavage was allowed to proceed for 1 h at 37 °C.

### CN-PAGE and in-gel fluorescence imaging

The cleaved samples were mixed with 4X native loading buffer (120 mM Tris pH 6.8, 20% glycerol, 0.02% bromophenol), and each sample was loaded onto an AnykD precast polyacrylamide gel (Bio-Rad, cat. no. 4569036). The gel was run in 25 mM Tris, 192 mM glycine, pH 8.3, at 70 V for 4 h at 4 °C. Protein LoBind tubes (Eppendorf, cat. no. 022431081) minimized potential adsorption of protein, and samples were protected from light throughout the entire process. The gel was then immediately imaged using an iBright 1500 imager (Invitrogen, excitation wavelength: 494 nm, emission wavelength: 512 nm).

### Preparation of the HR2 peptide library

To create the library for saturation mutagenesis of the HR2 peptide (shortHR2), the residues in the wildtype sequence were replaced by each of the 19 amino acids, one at a time. The 5’ and 3’ flanking sequences ‘GGTCAAAGCGGTCAAcatatg’ and ‘tctagaCGCATTAGTCCGATG’, respectively, were appended onto each coding DNA sequence. All these sequences were purchased as an oligonucleotide pool generated by massively parallel synthesis (Twist Bioscience, San Francisco, CA). The oligonucleotide sequences were amplified and gel purified. The 5’ end of the forward primer contained the sequence CATATG, the recognition motif for the NdeI restriction enzyme, while the 5’ end of the reverse primer contained the sequence TCTAGA, the recognition motif for the XbaI restriction enzyme. Both NdeI and XbaI recognition sites were introduced into the pBAD33 backbone containing eCPX. The oligo pool and the pBAD33 plasmid were then cut by NdeI/XbaI and ligated (using T4 DNA ligase) in the ligase buffer to circularize the linear product. We transformed the ligated DNA into NEB® 5-alpha Electrocompetent *E. coli*. We chose to perform the DMS in *E. coli* rather than a eukaryotic system (such as HEK293T cells) because this platform has been successfully used and validated in our previously published work^52^, ensuring a highly robust and reliable baseline for this study. To enable HR1 binding, the HR2 variant library was fused to the N-terminus of the bacterial surface-display scaffold, eCPX, and electroporated into *E. coli*. Electroporation enables the easy transformation of libraries containing millions of unique variants. Mammalian expression, by contrast, requires either transient transfection or stable integration. Because mammalian cells divide much more slowly than bacteria, scaling up culture volumes to achieve the same library coverage becomes incredibly resource- and time-intensive. Furthermore, maintaining a strict low multiplicity of infection/transfection is critical in mammalian cells to avoid expressing multiple variants per cell, which would otherwise lead to a false interpretation of the binding readout. In addition, the selection process—sorting the bound population via flow cytometry—is rapid in *E. coli*, resulting in a high-throughput workflow with a short turnaround time. Library coverage was maintained at ∼1,000-fold per variant and assessed by colony counts of transformants. We isolated and saved mini-prepped plasmid for use in the surface-display assay.

### Purification of HR1 for the surface display assay

HR1 complementary DNA (cDNA) was inserted between a His6-SUMO sequence and the GFP gene and cloned into a pSMT3 vector. The SUMO-HR1-GFP fusion was expressed in BL21(DE3) cells by overnight induction with 0.5 mM isopropyl β-D-1-thiogalactopyranoside at 18 °C. Cells were pelleted and then resuspended in a buffer containing 50 mM Tris, pH 8.0, 300 mM NaCl, 10 mM imidazole, 2 mM β-mercaptoethanol, and a mixture of protease inhibitors.

Cells were lysed by French press, and the lysates were clarified by centrifugation at 35,000 × g for 1 h. The SUMO-HR1-GFP fusion was purified using a HisTrap Fast Flow column (Sigma). Eluted fractions were then dialyzed against a buffer containing 20 mM Tris-HCl (pH 8.0), 150 mM NaCl, and 1 mM dithiothreitol at 4 °C overnight. After dialysis, the sample was concentrated and gel-filtered in the same buffer on a HiLoad 16/60 Superdex 200 (GE Healthcare) column.

Peak fractions that contain SUMO-HR1-GFP fusion were pooled and concentrated to 4 mg/mL through centrifugation with a 10 kDa MWCO concentrator (EMD Millipore cat. no. UFC901008).

### Surface display of the HR2 peptide library

We used 100 ng DNA (shortHR2 peptide library) to transform 100 μL electrocompetent *E. coli* MC1061 cells. Details for the peptide expression can be found in Shah *et al.*^53^. After induction of peptide expression on the N-terminus of the surface display scaffold eCPX^54^, *E. coli* cells were spun down and resuspended in a buffer containing 50 mM HEPES (pH 7.5), 150 mM NaCl, 0.2% bovine serum albumin, and incubated with 1 μM SUMO-HR1-GFP on ice for 1 h. The labeled and washed cells were then resuspended and diluted 5-fold in the same buffer for sorting. Cells were sorted on a Sony SH800 cell sorter. The distribution of GFP fluorescence (HR1 binding) for the whole cell population was measured, and a sorting gate was set to collect the highest 25–35% of this distribution. For each sample, 1,000,000 cells were collected into a 15 mL conical tube containing 3 mL of LB on ice. For each library, an analogous tube was prepared with approximately 1,000,000 unsorted cells diluted in LB. Details of the DNA isolation and deep sequencing can be found in Shah *et al.*^53^.

### Analysis of deep sequencing data

A custom shell script utilizing grep was used to count the number of occurrences of each variant in the raw sequencing file. Enrichment ranks were calculated following a previously described method^55^. Briefly, for each HR2 variant *i*, the enrichment rank (*E^i^*) was computed based on its relative abundance in the GFP+ sorted population compared to the unsorted population and normalized to the corresponding ratio for the wildtype (WT) HR2 sequence. The enrichment rank was calculated using the following formula:

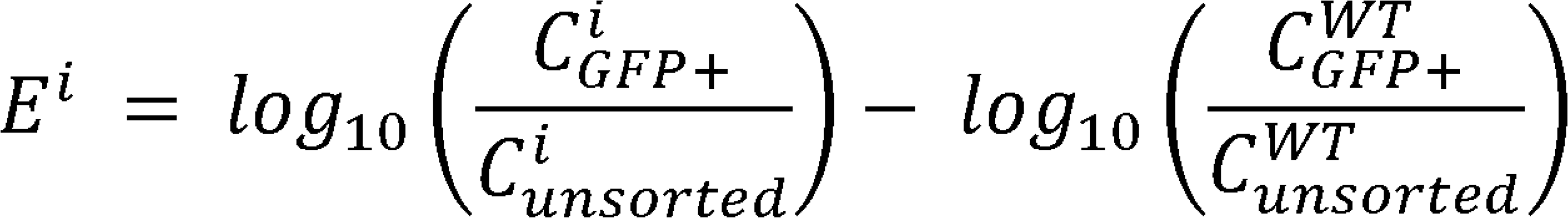

where *C^i^* and *C^WT^* refer to the counts of the HR2 variant *i* and the wildtype HR2, respectively, in either the GFP+ or unsorted populations. Using this approach, an enrichment rank of +1 or −1 corresponds to a 10-fold increase or decrease, respectively, in the abundance of a variant relative to wildtype in the GFP+ population compared to the unsorted population.

### Peptide synthesis and preparation

All the non-fluorescently labeled peptides listed in **Fig. 1a** were synthesized by GenScript USA Inc. HPLC and liquid chromatography–mass spectrometry profiles for the three new designs (HR2-QHN, HR2-NNS, and HR2-NHN) are shown in **Figs. S4 to S6** (provided by the manufacturer).

We developed a new protocol for dissolving HR2 peptides with the following goals: (A) minimize the contact of peptides with other substances, such as DMSO and concentrator membranes, that may introduce contaminants that interfere with concentration calculations by measuring absorbance at 205 nm; (B) minimize the leftover salts and other substances from the peptide powder, such as trifluoroacetic acid, that may be harmful to cells and viruses; (C) keep the protocol simple enough that it can be easily reproduced in most biology laboratories. A 5-mL 1800-Da-MWCO desalting column (Thermo Scientific, cat. no. 43426) was prepared according to the manufacturer’s manual and equilibrated with 10 mL of 20 mM HEPES-Na pH 7.5. 2 mg of the lyophilized peptide powder from GenScript was resuspended in 0.5 mL ddH_2_O. To clarify the peptide suspension, 0.1 M NaOH was added dropwise until it turned clear. The clarified peptide solution was immediately applied to the pre-equilibrated column and allowed to enter the column bed. 1 mL aliquots of 20 mM HEPES-Na pH 7.5 were applied to the column sequentially, and fractions were collected in 1.5-mL Eppendorf Protein LoBind tubes (Eppendorf, cat. no. 022431081). The peptide concentration for each fraction was quantified with a Nanodrop spectrometer by measuring absorbance at 205 nm. Usually, the peptide should be in fraction #2 with a concentration of 0.5–0.8 mg/mL, or around 200 μM.

Detailed comparisons for preparing peptides in solution using different protocols are rarely published, so we only compared our new protocol using a desalting column with our previously published protocol using a concentrator^6^. Using our cell-cell fusion assay, we found that the IC_50_ barely changes for the well-behaved longHR2_42 (**Fig. S11**). However, the IC_50_ for the shortHR2 decreased from 263 nM to 27 nM (**Fig. S11**), indicating that the new protocol either improves the sample purity substantially, or only removes trace amounts of contaminants that may have substantial impact on concentration measurement by absorption at 205 nm^56^. The most commonly used way of protein concentration measurement by absorption at 280 nm is not applicable since the HR2 peptides do not have any aromatic residues, while accurate quantification by mass of peptide powder requires an overwhelmingly large amount than needed in our screening assay, plus the weight of peptide powder inevitably contains that from salts during lyophilization.

### HEK cell-cell fusion assay

We optimized the cell-cell fusion assay based on the α-complementation of *E. coli* β-galactosidase for comparing the inhibitory activity of different peptides with higher throughput. Suspension culture Expi293F cells (Thermo Fisher, cat. no. A14527), a clonal derivative of Human Embryonic Kidney (HEK) 293 cells, were grown to a density of 1–2 × 106 cells/mL in FreeStyle 293 expression medium (Thermo Fisher, cat. no. 12338026). The cells were then pelleted, resuspended in medium to a density of 1 × 10^6^ cells/mL, and allowed to recover at 37 °C for 30 min. One group of cells was then co-transfected using polyethyleneimine (PEI, Sigma) (125 μg PEI/mL cells) with the full-length SARS-CoV-2 Wuhan-Hu-1 strain S protein construct (12.5 μg DNA/mL cells) and the α-fragment of *E. coli* β-galactosidase construct (12.5 μg DNA/mL cells) to generate the S protein-expressing cells. Since HEK cells do not express TMPRSS2, it is likely that S is cleaved by other host cell proteases that render S fusogenic upon binding to its receptor ACE2. Using the same amount of PEI, the other group of cells was co-transfected with the full-length ACE2 (12.5 μg DNA/mL cells) construct and the ω-fragment of *E. coli* β-galactosidase construct (12.5 μg DNA/mL cells) to generate the ACE2 receptor-expressing cells. Note that the endogenous ACE2 level is very low in HEK cells, requiring ectopic expression for efficient fusion. As a negative control, two additional groups of cells were transfected with either the α-fragment or the ω-fragment of *E. coli* β-galactosidase construct alone. After incubation of the cells in flasks at 37 °C for 24 h, the cells were pelleted. The S-expressing cells were resuspended in FreeStyle 293 expression medium supplemented with different concentrations of peptide or peptide-TMPRSS2 mixture (50 μL, 2 × 10^6^ cells/mL), respectively. The ACE2-expressing cells and negative control cells were resuspended in 50 μL fresh FreeStyle 293 expression medium (pH 7.4) to be 2 × 10^6^ cells/mL. S-expressing, and ACE2 cells or α-fragment and ω-fragment cells were then mixed in a 96-well plate (Greiner Bio-One) to initiate cell-cell fusion at 37 °C for 2 h. Fusion was arrested by adding 100 μL β-galactosidase substrate from the Gal-Screen reporter system (Invitrogen). The mixture was incubated at 37 °C in the dark for 1 h before luminescence was recorded using a Tecan Infinite M1000.

### Cryo-EM structure determination

The cryo-EM structure of the HR2-NHN bound to HR1 was determined following a molecular scaffolding method described previously^41^. Briefly, the scaffolded HR1/HR2 complex was generated by co-expressing the scaffolded HR1 and Small Ubiquitin-like Modifier (SUMO)-tagged HR2-NHN in *E. coli* BL21(DE3) using auto-inducing lysogeny broth (LB) medium^51^, followed by nickel affinity chromatography and size-exclusion chromatography (SEC) with a Superose 6 Increase 10/300 GL column in 25 mM Hepes-Na, pH 7.4, 150 mM NaCl, 0.5 mM EDTA, 0.5 mM Tris(2-carboxyethyl)phosphine (TCEP). The sample was concentrated to 20 μM, supplemented with 0.05% Nonidet P-40, and plunge frozen on a Quantifoil 2/1 holey carbon grid using a Vitrobot Mark IV (Thermo Fisher). A total of 20,031 movies were recorded using a Titan Krios electron microscope (Thermo Fisher) equipped with a K3 camera (Gatan) using the EPU automation software (Thermo Fisher Scientific), at a nominal magnification of 130,000× and a pixel size of 0.325 Å. Each movie contained 40 frames with a total electron dose of 52 e^−^/Å^2^. The data were processed using a combination of MotionCor2^57^, Gctf^58^, EMAN2^59^, cryoSPARC^60^, and RELION^61^, as described previously^41^. More details for data collection and processing are summarized in **Fig. S7 and Table S1.**

For model building, the PDB 8czi was used as the template. The mutations of HR2 were performed in Coot^62^ and then refined by the automated structure refinement protocol in Rosetta^63^. The structure was then subjected to real-space refinement (minimization_global, local_grid_search, adp) in PHENIX^64^. Coot^62^ was used for further fitting of sidechains and manual inspection.

### Animals

A total of 78 female BALB/c mice (Envigo, Indianapolis, IN, USA) were transferred to the University of Helsinki biosafety level 3 (BSL-3) facility and acclimatized to individually ventilated biocontainment cages (ISOcage; Scanbur, Karlslunde, Denmark) for 7 days with ad libitum water and food (rodent pellets). For subsequent experimental infection, the mice were placed under isoflurane anesthesia and inoculated intranasally with 20 μL of virus dilution or DMEM (non-infected control). The animals were held in an upright position for a few seconds to allow the liquid to flow downwards in the nasal cavity. Mice did not lose weight when infected with the SARS-CoV-2 Beta variant. Their well-being was further monitored carefully for signs of illness (e.g., changes in posture or behavior, rough coat, apathy, ataxia). Euthanasia was performed under terminal isoflurane anesthesia with cervical dislocation. Experimental procedures were approved by the Animal Experimental Board of Finland (license number ESAVI/32448/2023).

### Cells

A549 stably expressing human *ACE2* and *TMPRSS2* fused to GFP (A549-AT)^45^ were grown in Dulbecco’s modified Eagle’s medium (DMEM), 2 mM glutamine, 10% heat-inactivated fetal bovine serum (FBS), and 1X penicillin/streptomycin.

### Virus isolation, propagation, and sequencing

All experiments with wildtype or mutant SARS-CoV-2 were performed in BSL-3 facilities of the University of Helsinki with appropriate institutional permits. Viruses isolated from consenting COVID-19 patients (permits HUS/32/2018 § 16) were propagated once in VeroE6-TMPRSS2 cells^43^, titrated by plaque assay, and stored at −80 °C in DMEM, 2% FBS, and 1X penicillin/streptomycin. All SARS-CoV-2 viruses were sequenced by next-generation sequencing at the University of Helsinki. The SARS-CoV-2 Beta variant virus used in mice experiments was isolated from infected patient nasopharyngeal samples as described^43^ and amplified using VeroE6-TMPRSS2^43^. Once propagated, their genomic sequence was confirmed using an Illumina platform available at the Department of Virology, University of Helsinki^43^. The Beta variant sequence has been previously described in detail^46^ and deposited in the NCBI GenBank database under accession number MW717678. For virus production, at 3 days post inoculation, the collected medium from the infected cells was centrifuged twice at 4,500 × g for 10 min at 4 °C, and the cleared supernatant was aliquoted in cryotubes and stored at −80 °C in the BSL-3 facility.

The virus was propagated in Minimum Essential Medium (MEM) containing 2% FBS, 20 mM HEPES, pH 7.2, 2 mM glutamine, and 1X penicillin/streptomycin. Virus titrations were performed by standard plaque assay in VeroE6-TMPRSS2 cells as previously described^43^. Infected cells were maintained in incubators at 37 °C and 5% CO_2_ in the BSL-3 facility at Helsinki University Hospital.

### Virus infection and peptide treatments in cell cultures

Human bronchial adenocarcinoma-derived lung Calu-3 cells (ATCC, cat. no. HTB-55; male) and A549-AT^65^ cells were seeded in DMEM containing 10% (A549-AT) or 20% (Calu-3) Dulbecco’s Modified Eagle’s High Glucose, pyruvate Medium (DMEM, Gibco, cat. no. 11995065), 1X non-essential amino acids (NEAA, Gibco, cat. no. 11140050), 10% heat-inactivated fetal bovine serum (FBS; Gibco, cat. no. 26140079), and 100 U/mL penicillin/streptomycin (Gibco, cat. no. 15140122). A549-AT cells were seeded at 15,000 cells per well and Calu-3 cells were seeded at 22,000 or 12,000 cells per well in 96-well imaging plates (cat. no. 6005182; PerkinElmer) 24 h (for time 0 hbi experiments) or 96 h (for the 72 hbi experiments) before infection with SARS-CoV-2 mCherry^66^ (a recombinant virus based on the original Wuhan-Hu-1 strain) or the Omicron XBB.1.5 variant. The peptides were diluted in the same infection medium and added at the indicated times. The amount of virus used to infect the cells was adjusted to obtain 15–20% infected cells in vehicle-treated controls, as determined by high-content fluorescence imaging at 18 hpi and automated image analysis after fixation with 4% paraformaldehyde (in PBS) for 20 min at room temperature.

### High-content fluorescence imaging

Fixed cells were washed three times with Dulbecco-modified PBS containing 0.2% BSA (DPBS/BSA), permeabilized with 0.1% Triton X-100 in DPBS/BSA, and nuclear DNA stained with Hoechst DNA dye (ThermoFisher Scientific, cat. no. H3570). Automated fluorescence imaging was performed using a Molecular Devices Image-Xpress Nano high-content epifluorescence microscope equipped with a 10× objective and a 4.7-megapixel CMOS camera (pixel size: 0.332 μm). Image analysis was performed with CellProfiler-4 software (www.cellprofiler.org). Nuclei were automatically detected using the Otsu algorithm included in the software. To automatically identify infected cells, an area surrounding each nucleus (5-pixel expansion of the nuclear area) was used to estimate the fluorescence intensity of the virally expressed mCherry fluorescent protein, using an intensity threshold such that <0.01% of ‘positive cells’ were detected in non-infected wells.

### Infections and peptide treatments in mice

Eight-week-old female BALB/c mice were anesthetized using isoflurane and intranasally inoculated (n = 4 or 5 per group) with 2 × 10^5^ PFU of the SARS-CoV-2 Beta variant in 20 μL DMEM^46^. The peptides solubilized as described above in 20 mM HEPES pH 7.2 at a concentration of 1 mg/mL (or 0.1 mg/mL when indicated) were applied as a 20 μL drop/mouse at indicated times. The virus and peptide drops were applied to the openings of the animal’s nostrils, and the liquid was naturally inhaled during anesthesia. At 48 hpi, animals were euthanized under terminal isoflurane anesthesia with cervical dislocation and dissected immediately after sacrifice. The right lungs were dissected, collected, and frozen at −80 °C before processing for PCR and analysis of viral RNAs, or for homogenization and virus extraction, as described in the next paragraph; left lungs were fixed in 10% buffered formalin for 48 h and stored in 70% ethanol for histological and immunohistochemical examinations.

### Lung homogenization and virus reinfection in VeroE6 cells

A portion of the right lung from each mouse was washed once with ice cold PBS, weighed, and 100 mg homogenized by mixing with 100 mg of glass beads (1.0 mm diameter, Sigma-Aldrich, cat. no. Z250473) in 500 μL ice-cold MEM containing 2% FBS, 20 mM HEPES pH 7.2, 2 mM glutamine, and 1X penicillin/streptomycin. Samples were vortexed three times at full speed for 30 s (total vortexing time 1.5 min). After centrifugation at 400 × g for 10 min, 25 μL of the supernatant from each sample was used to infect VeroE6 cells grown on 96-well imaging plates in a total volume of 100 μL. At 18 hpi cells were fixed in 4% PFA for 20 min at room temperature and processed for immunofluorescence staining of viral protein N, automated imaging and image analysis as described above.

### RNA Isolation and qRT-PCR

RNA was extracted from lung samples using Trizol (Thermo Scientific) according to the manufacturer’s instructions. Isolated RNA was subjected to one-step qRT-PCR analysis as described using primers specific for the viral genome encoding for the subgenomic E gene^67^ with TaqMan fast virus 1-step master mix (ThermoFisher Scientific, cat. no. 4444432) using AriaMx instrumentation (Agilent, Santa Clara, CA, USA). The actin qRT-PCR is described in Zivcec *et al.*^68^.

Primer and probe sequences used in the qRT-PCR.

Subgenomic E Forward cgatctcttgtagatctgttctc^67^

Probe acactagccatccttactgcgcttcg^67^

Reverse atattgcagcagtacgcacaca^67^

Beta-actin Forward actgccgcatcctcttcct^68^

Probe cctggagaagagctatgagctgcctgatg^68^

Reverse tcgttgccaatggtgatgac^68^

### Immunohistochemistry

The left lungs of the sacrificed mice were trimmed for histological examination and paraffin-wax embedded. The heads were sawn longitudinally in the midline using a diamond saw (Exakt 300; Exakt, Oklahoma, OK, USA), then decalcified and processed as previously described^46^. Consecutive sections (3–5 μm) were prepared from lungs and subjected to immunohistochemistry for the detection of SARS-CoV-2 antigen expression using a rabbit polyclonal anti-SARS-CoV NP antibody that cross-reacts with NP of SARS-CoV-2 (Rockland Immunochemicals, Limerick, USA, cat. no. 200-402-A50).

### Statistics and data analysis

Data from the HEK cell-cell fusion assay from three independent biological replicates, determined for each concentration of inhibitor, are used. For the HEK cell-cell fusion assay, normalized fusion was calculated as (Luminescence_(+inhibitor)_ − Luminescence_(α&ω)_)/(Luminescence_(+PBS)_ − Luminescence_(α&ω)_), where “+inhibitor” or “+PBS” refers to adding inhibitor or PBS to the mixture of the cells expressing the α-fragment of *E. coli* β-galactosidase and S, and the cells expressing the ω-fragment of *E. coli* β-galactosidase and ACE2, and “α&ω” refers to the mixture of the cells expressing the α-fragment only and the cells expressing the ω-fragment only.

After the normalization, the arithmetic means of the three replicates were used to fit the inhibition curves to obtain estimates of the IC_50_ values; the estimates were obtained by nonlinear regression of inhibitor concentration vs. response in GraphPad Prism version 9.1.0 for macOS (GraphPad Software, San Diego, CA, https://www.graphpad.com). The fitted model is Y = Bottom + (Top − Bottom)/(1 + (IC_50_/X)^HillSlope^), where Y is the extent of inhibition, X is the inhibitor concentration, Bottom and Top are the minimal and maximal inhibition. The SE of the IC_50_ estimation was calculated using OriginPro 9.1 (OriginLab Corporation).

For the peptide dose responses *in vitro* in Calu-3 cells (**Fig. 5 and Fig. S8**), and for the estimation of bioavailability in mice (**Fig. 6 and Fig. S9**), all statistical analyses were performed in R. The R code used is available in the GitHub repository (available at the link: https://github.com/bongrita/SARSCoV_Fusion_inhibitor_peptide.git).

For the *in vitro* dose-response experiments (**Fig. 5**), normalized infections were modeled as a function of peptide concentration using four-parameter log-logistic regression (LL.4, drc package of R)^69^. The IC_50_ values were defined as the concentration at which normalized infection reached a value of 0.5 and were estimated using the “ED()” function of R with 95% confidence intervals calculated by the delta method.

IC_50_ comparisons for longHR2_42 and HR2-NHN were performed using joint dose-response models and the R function “compParm()” ratio test^69^. To test if the difference between the fitted curves was statistically significant, area under the fitted curve (AUC) was calculated over 0.1– 1,000 nM peptide concentrations by trapezoidal integration and compared between the two peptides using non-parametric bootstrap resampling with 1,000 iterations^70^. AUC was recalculated for each iteration, with 95% confidence intervals derived from the bootstrap percentile distribution and two-sided P values calculated from the bootstrap tail probabilities.

For the *in vivo* mouse re-infection time-course experiment, infection measurements from vehicle- or peptide-treated mice were normalized to the mean infection level of the vehicle-treated control group, which was defined as 1.0, and data are presented as mean ± SD. Overall infection across the 0–24 h pretreatment interval was summarized by trapezoidal AUC calculated from the observed mean normalized infection values at 0, 4, 8, 16 and 24 h, and differences between peptides were assessed by bootstrap resampling of mice within each time point with 1,000 iterations^70^. To estimate the duration of antiviral activity (T_50_), a logistic time-response curve was fitted separately for each peptide using data from 0–16 h; the 24-h time point was excluded from T_50_ estimation because the decrease between 16 h and 24 h violated the monotonicity assumption of the logistic model. T_50_ was defined as the pretreatment interval at which fitted normalized infection reached 0.5, corresponding to 50% of the vehicle-treated infection level. Differences in the AUC for each peptide were assessed by bootstrap resampling within each time point with 1,000 iterations^70^. For all analyses, statistical significance was defined as P < 0.05.

Statistical analysis of the mouse experiment described in **Fig. 6b** was performed with Ordinary one-way Anova Dunnett’s multiple comparison test using GraphPad Prism version 9.1.0 for macOS (GraphPad Software, San Diego, CA, https://www.graphpad.com). For the mouse treatment “0 h 0.1 mg/kg”, two of the mice died during anesthesia prior to infection and thus statistical analysis was not performed with the remaining 2 mice.

### Figure preparation

The figures of PDB structures and maps were made using UCSF Chimera^71^. The data fitting of all inhibition assays was performed and plotted using GraphPad Prism version 9.1.0 for macOS (GraphPad Software, San Diego, CA, https://www.graphpad.com) or R. Figure panels were assembled with Adobe Illustrator and Image-J (Fiji).

### Data availability

The EM maps and corresponding structures reported here have been deposited in EMDB (Electron Microscopy Data Bank) and PDB with the following accession IDs: EMD-70569 and PDB 9okn. Authors will release the atomic coordinates and experimental data upon article publication. The R code used for statistical analysis, generation of graphs, and calculation of AUCs and IC_50_s is available on GitHub (accessible at the link: https://github.com/bongrita/SARSCoV_Fusion_inhibitor_peptide.git).

## Acknowledgments

We thank Bing Chen for kindly providing plasmids and protocols for the cell-cell fusion assay, Rebekah C Gullberg, Casey Thompson, Yuhao Li, Sean Whelan, and Judith Frydman for stimulating discussions, the Sarafan ChEM-H High-Throughput Screening Knowledge Center for providing the Tecan microplate reader, Robert Sullivan (University of Queensland) for the help in the histology of mouse lungs, and the Finnish Institute for Molecular Medicine (FIMM) High Content Imaging unit for providing imaging services. We also thank the Stanford Cryo-Electron Microscopy Center (cEMc) and the Stanford-SLAC Cryo-EM Center (S^2^C^2^) for support. This article is subject to HHMI’s Open Access to Publications policy. HHMI laboratory heads have previously granted a nonexclusive CC BY 4.0 license to the public and a sublicensable license to HHMI in their research articles. Pursuant to those licenses, the accepted manuscript of this article can be made freely available under a CC BY 4.0 license immediately upon publication.

## Author contributions

Conceptualization: KY, JK, GB, and ATB

Molecular cloning, protein expression, purification, and labeling: KY, KIW, RAP, and LE TMPRSS2 cleavage test: KY

Deep mutational scanning: SMur, SS, and JK Design of mutant peptides: KY

Cell-cell fusion assay: CW Structure determination: KY

Virus-Cell and -mouse infection assays: FT, RK, LK, SMak, SHS, TStr, TB, YTB, TSir, JH, OV, MJ, and GB

Supervision: JK, GB, and ATB

Writing: KY, CW, SMur, RK, KIW, JK, GB, and ATB

All authors have read and approved the article.

