## Supplementary material for "A rational design strategy and validation for protease-resistant fusion-inhibitor antiviral peptides": SI

**This PDF file includes:**

Figs. S1 to S11

Table S1

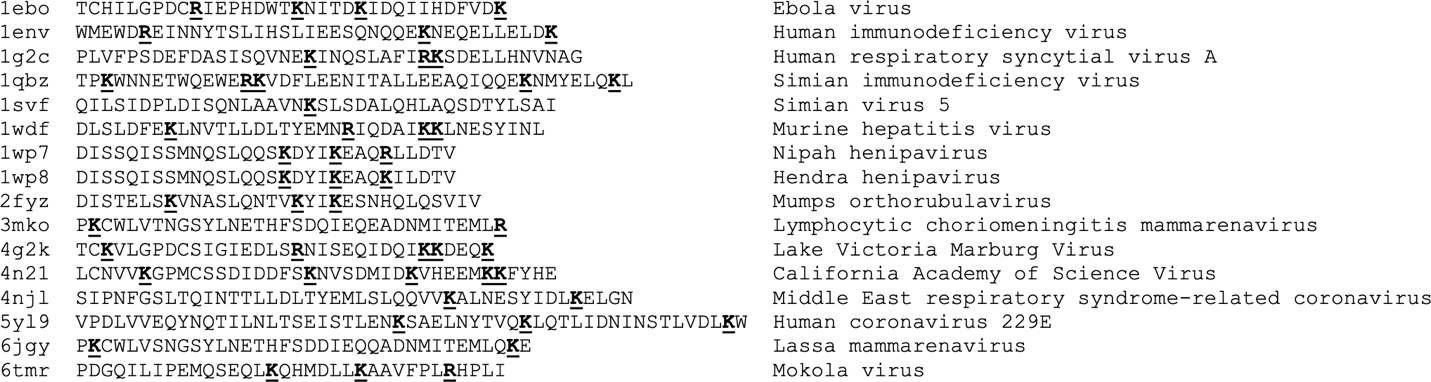

**Fig. S1. Basic residues are common in HR2-equivalent regions of type I enveloped viruses.**

We surveyed the PDB for X-ray crystal structures of post-fusion bundles of fusion proteins from various type I enveloped viruses, and then listed the PDB code, the sequence of the HR2-equivalent region with basic residues highlighted, and the name of the virus, as shown in the figure.

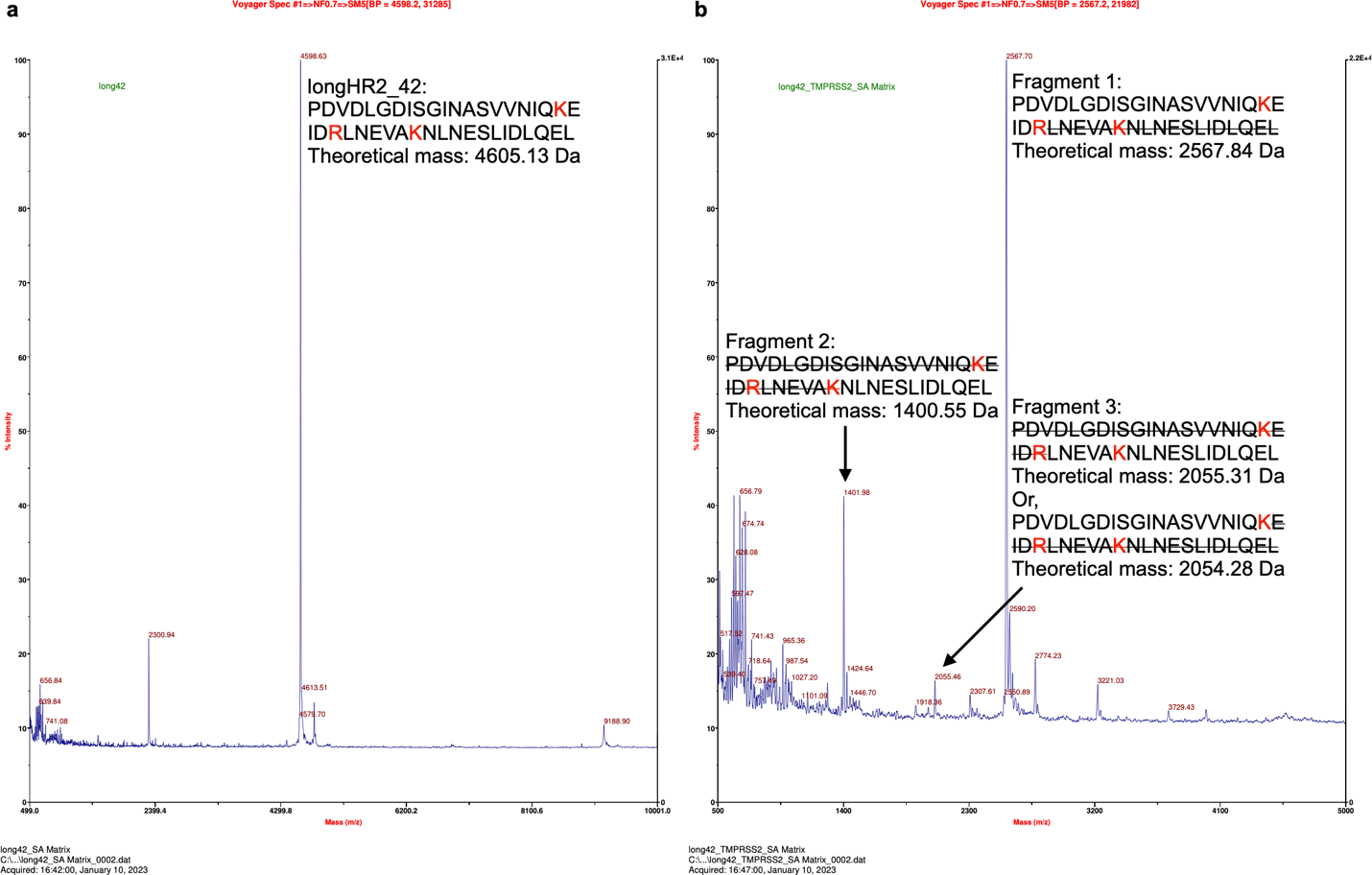

**Fig. S2. Mass spectrometry of longHR2_42 shows that TMPRSS2 cleaves the peptide at R1185 and K1191, and possibly at K1181.**

(**a**) Mass spectrometry of longHR2_42 before TMPRSS2 cleavage. The sequence and theoretical mass of intact longHR2_42 are indicated. All basic residues (K1181, R1185, and K1191) are colored red.

(**b**) Mass spectrometry of longHR2_42 after TMPRSS2 cleavage. The sequences and theoretical masses of cleaved fragments are indicated near their corresponding peaks. Sequences not present in the fragments are shown in strikethrough format for easier comparison with the intact sequence. Fragment 1 identifies R1185 as a cleavage site. Fragment 2 identifies K1191 as a cleavage site. Fragment 3, with a measured mass of 2055.46 Da, could correspond to two possible fragments with a theoretical mass of 2055.31 Da (cleavage at R1185) and 2054.28 Da (cleavage at K1181), respectively.

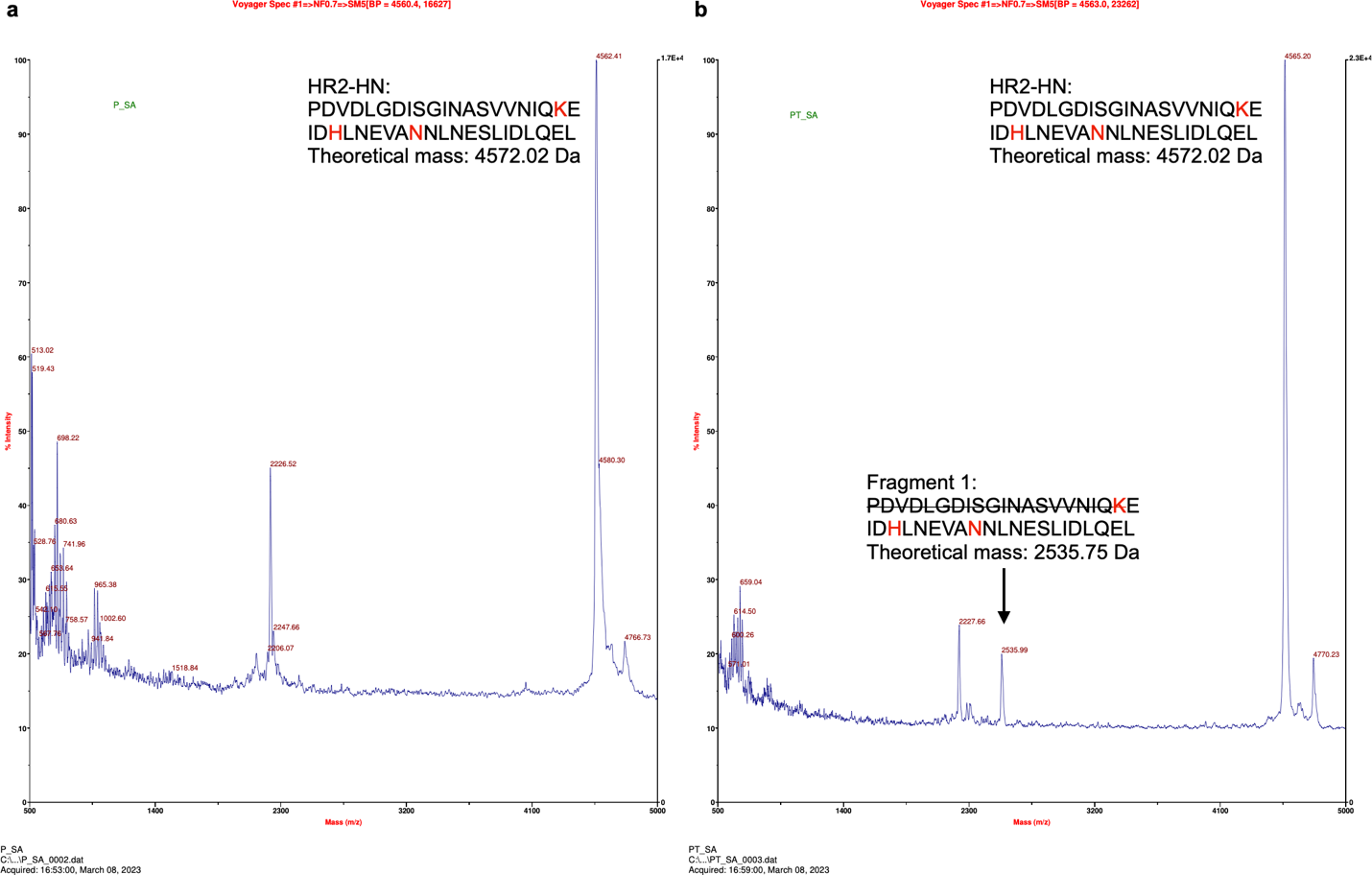

**Fig. S3. Mass spectrometry of HR2-HN shows that the TMPRSS2 cleavage site at K1181 is critical.**

We synthesized a mutant of HR2, named HR2-HN, with only the two confirmed cleavage sites mutated (R1185H and K1191N). We found that TMPRSS2 cleaves HR2-HN at K1181, albeit much less efficiently than it does the wildtype HR2 peptide (longHR2_42).

(**a**) Mass spectrometry of HR2-HN before TMPRSS2 cleavage. The sequence and theoretical mass of intact HR2-HN are indicated.

(**b**) Mass spectrometry of HR2-HN after TMPRSS2 cleavage. The sequences and theoretical masses of cleaved fragments are indicated near their corresponding peaks. Sequences not present in the fragments are shown in strikethrough format for easier comparison with the intact sequence.

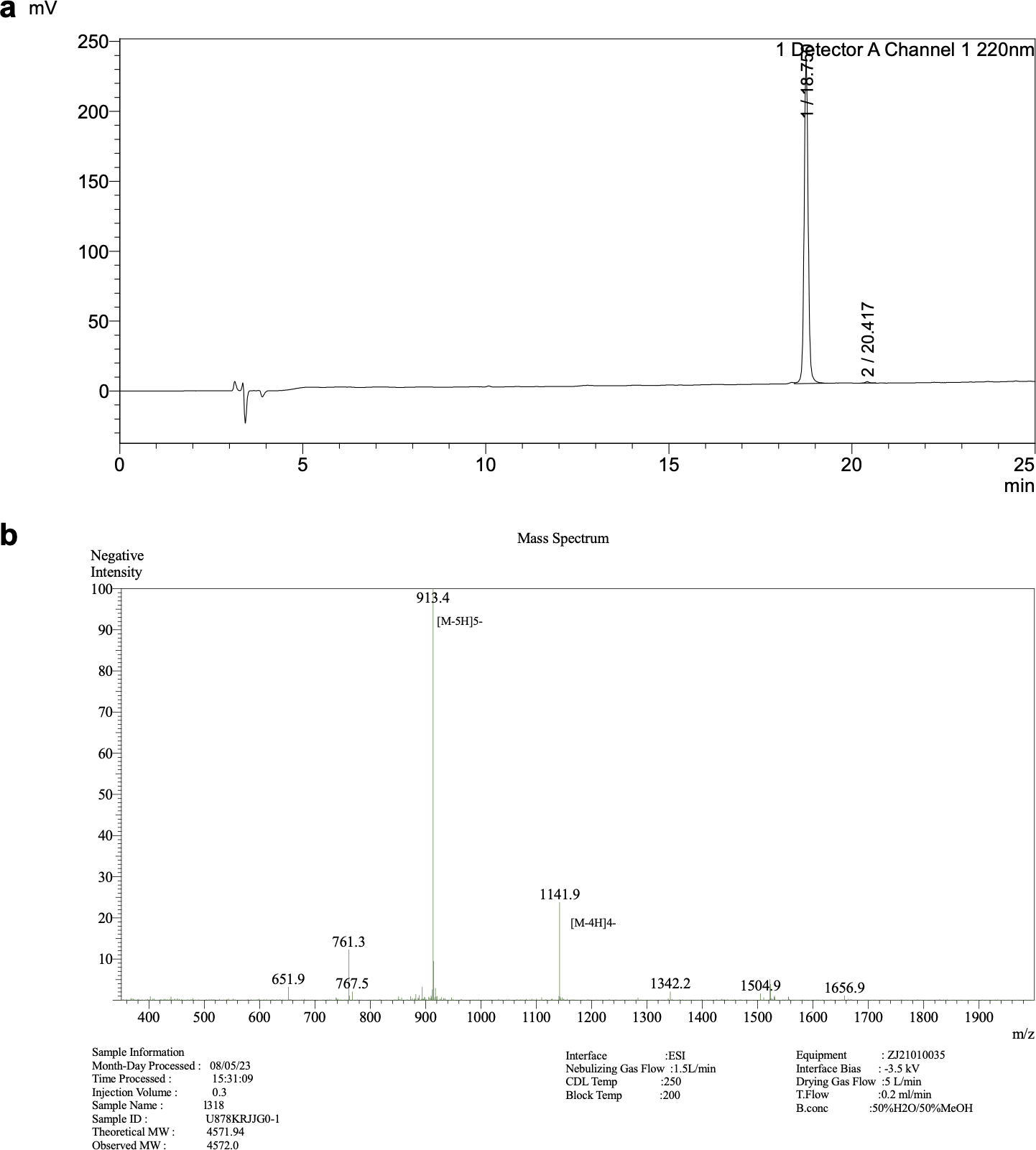

**Fig. S4. HPLC (a) and LC-MS (b) profiles of HR2-QHN.**

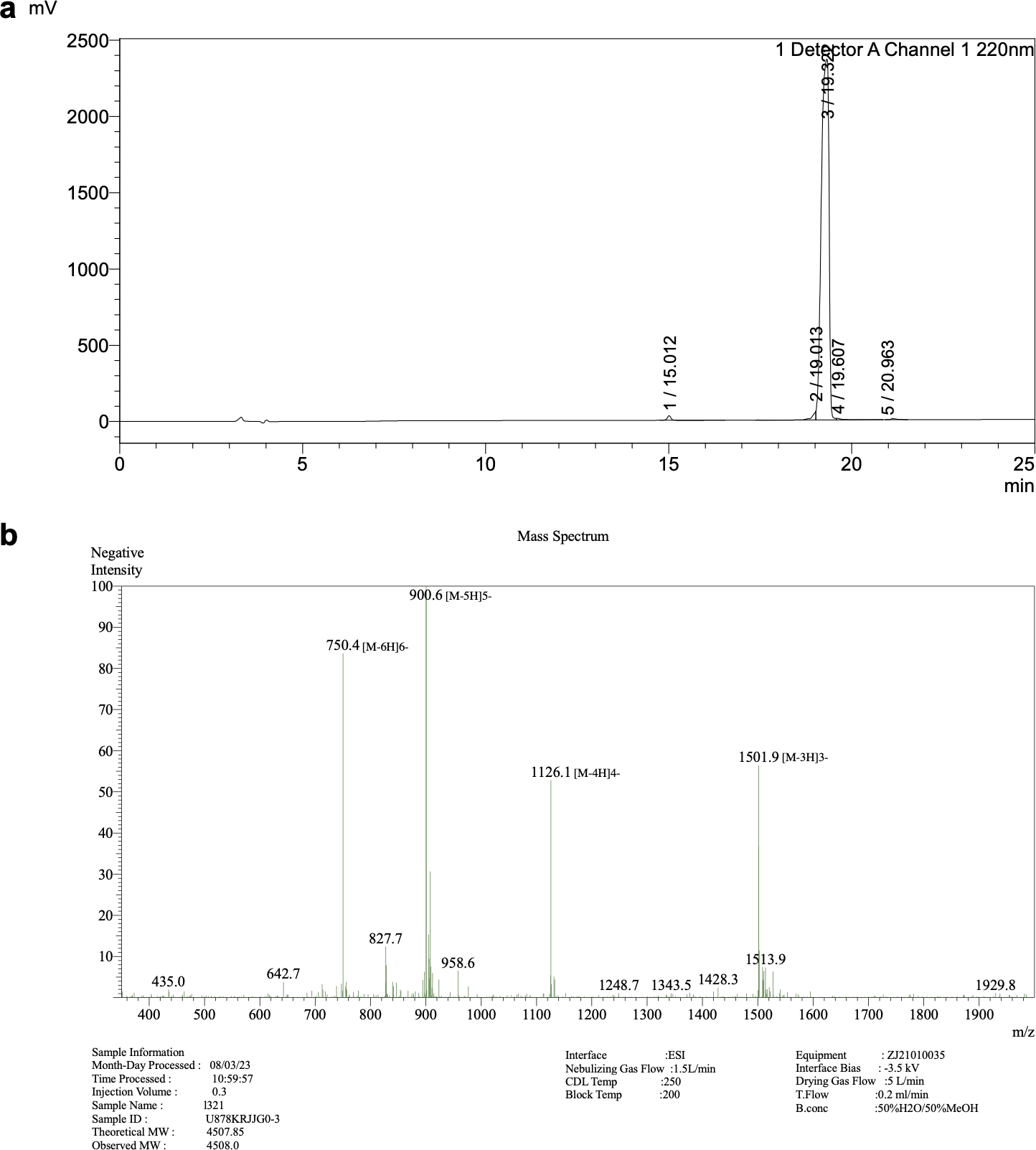

**Fig. S5. HPLC (a) and LC-MS (b) profiles of HR2-NNS.**

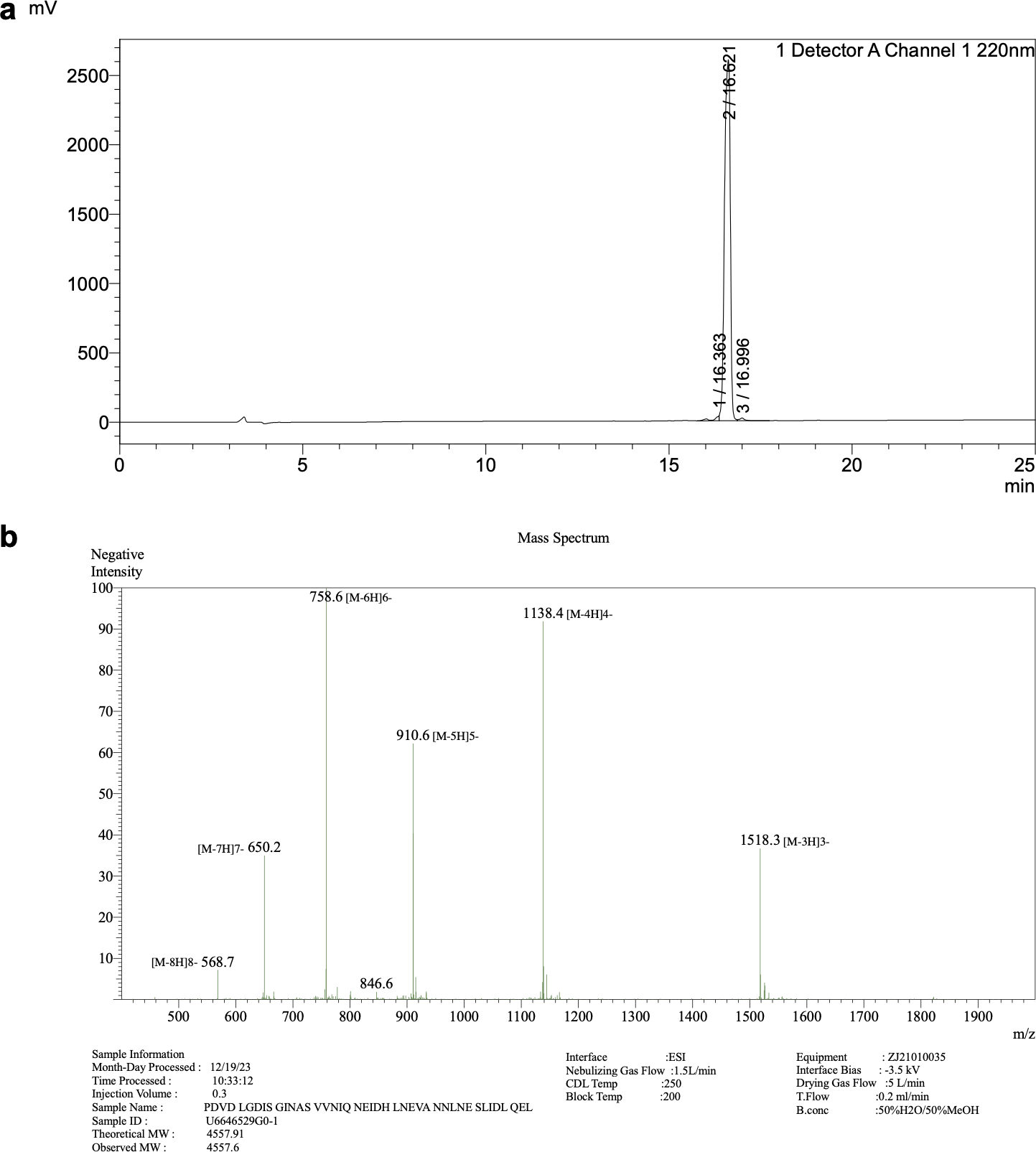

**Fig. S6. HPLC (a) and LC-MS (b) profiles of HR2-NHN.**

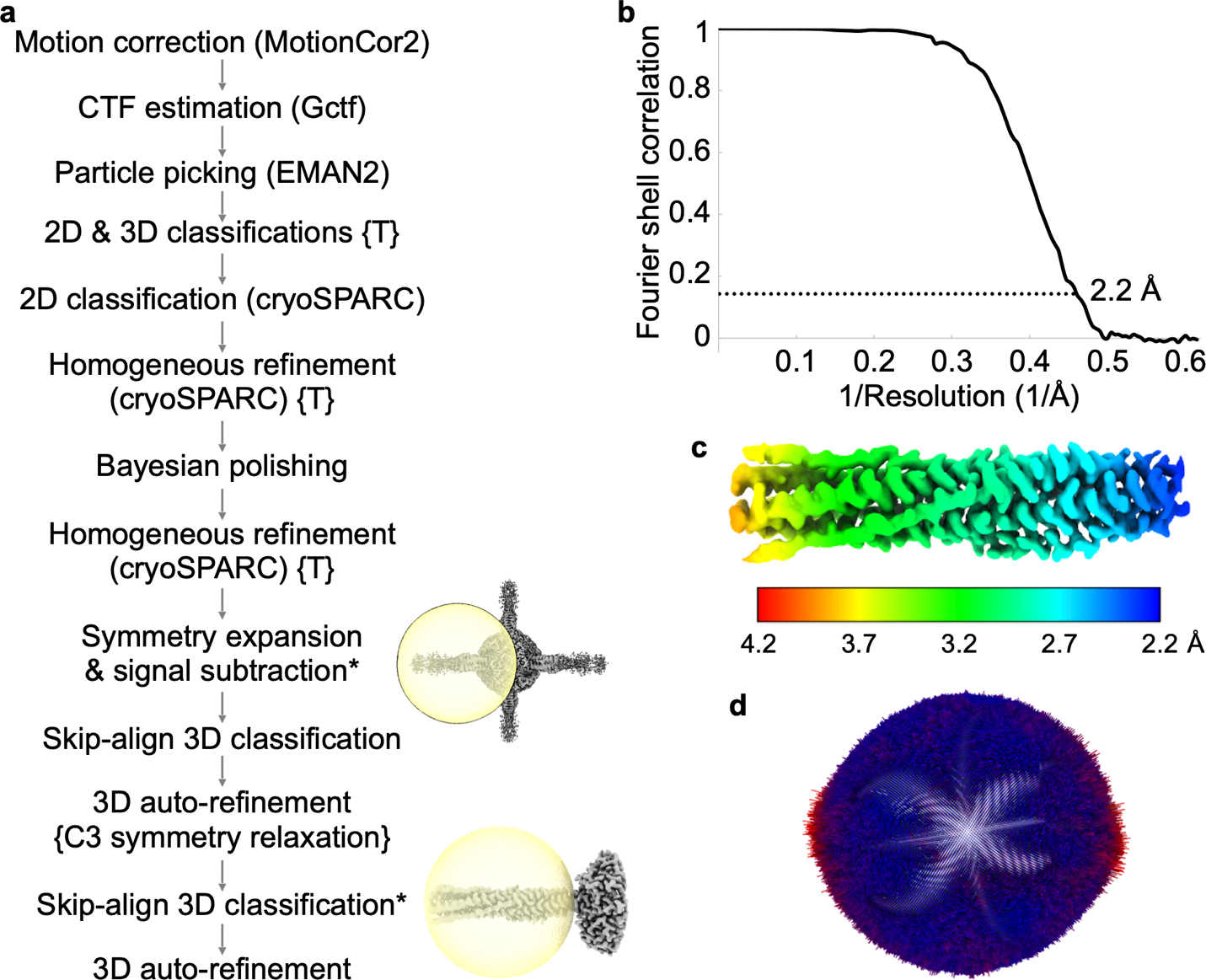

**Fig. S7. Cryo-EM structure determination of HR2-NHN bound to HR1.**

(**a**) Workflow of cryo-EM data processing. RELION was used for each step unless another program is indicated in parentheses. Symmetry was imposed or relaxed as indicated in curly brackets. The steps that used a manually generated mask (rather than the default spherical mask in RELION or the default dynamic masks in cryoSPARC) are indicated with a star sign and an image showing the manually generated mask (yellow) and an average map (gray).

(**b**) Fourier shell correlations (FSC) of the final local refinements with RELION (last 3D auto-refinement). Note that the FSC calculation was performed using the default spherical mask that covers both the HR1/HR2 bundle and part of the scaffold.

(**c**) Local resolution of the final reconstruction.

(**d**) Distribution of the particles’ orientations in the final reconstruction depicted in the same orientation as the map in panel **c**. The length of each bar is proportional to the number of particles oriented in that direction. The bars are also colored by length, with red indicating more particles and blue fewer.

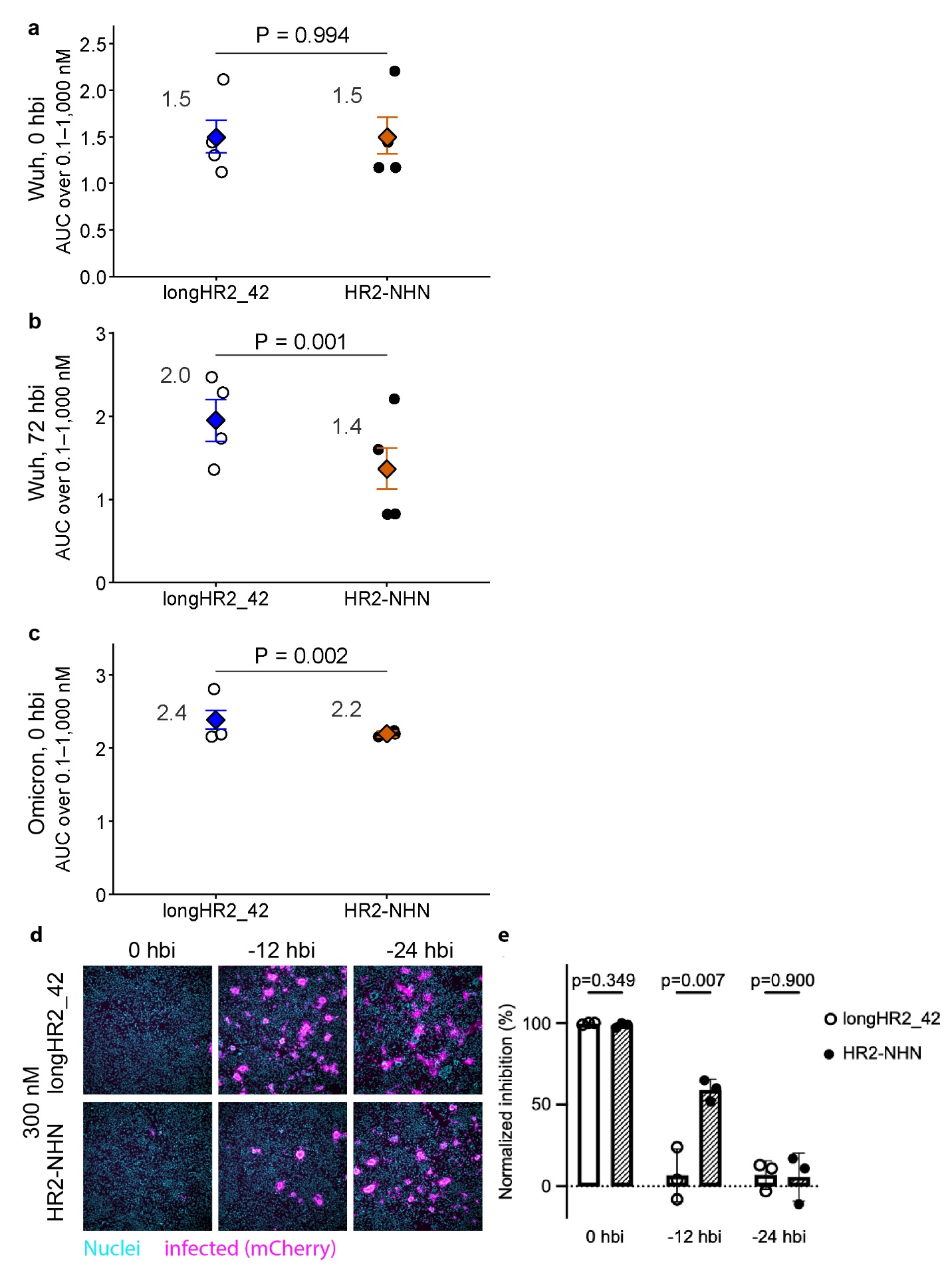

**Fig. S8. Comparison between longHR2_42 and HR2-NHN inhibitory activities when preincubated with cells for different times before infection.** Comparison of dose-response AUC between longHR2_42 and HR2-NHN when administered to Calu-3 cells at 0 hbi (**a**) and 72 hbi (**b**) with SARS-CoV-2-mCherry Wuhan-Hu-1, and at 0 hbi with Omicron XBB.1.5 (**c**). AUC was calculated over a peptide concentration range of 0.1–1,000 nM. Individual points represent AUC values calculated from independently repeated dose-response series. Diamonds indicate the pooled AUC estimate, and error bars indicate the bootstrap 95% confidence interval. Exact bootstrap P values for the comparison between peptides are shown. Blue, longHR2_42; orange, HR2-NHN. (**d**) Example cell images and (**e**) normalized inhibition of infection of A549-AT cells by recombinant SARS-CoV-2-mCherry in the presence of 300 nM longHR2_42 or HR2-NHN administered 0, 12, or 24 h before infection. Nuclei are stained with Hoechst DNA dye (cyan), and infected cells are detected with mCherry red fluorescent protein (magenta) expressed by the recombinant virus. The P values indicating the statistical significance of each sample pair are indicated in the graph. Statistical analysis was performed by the unpaired two-tailed t-test.

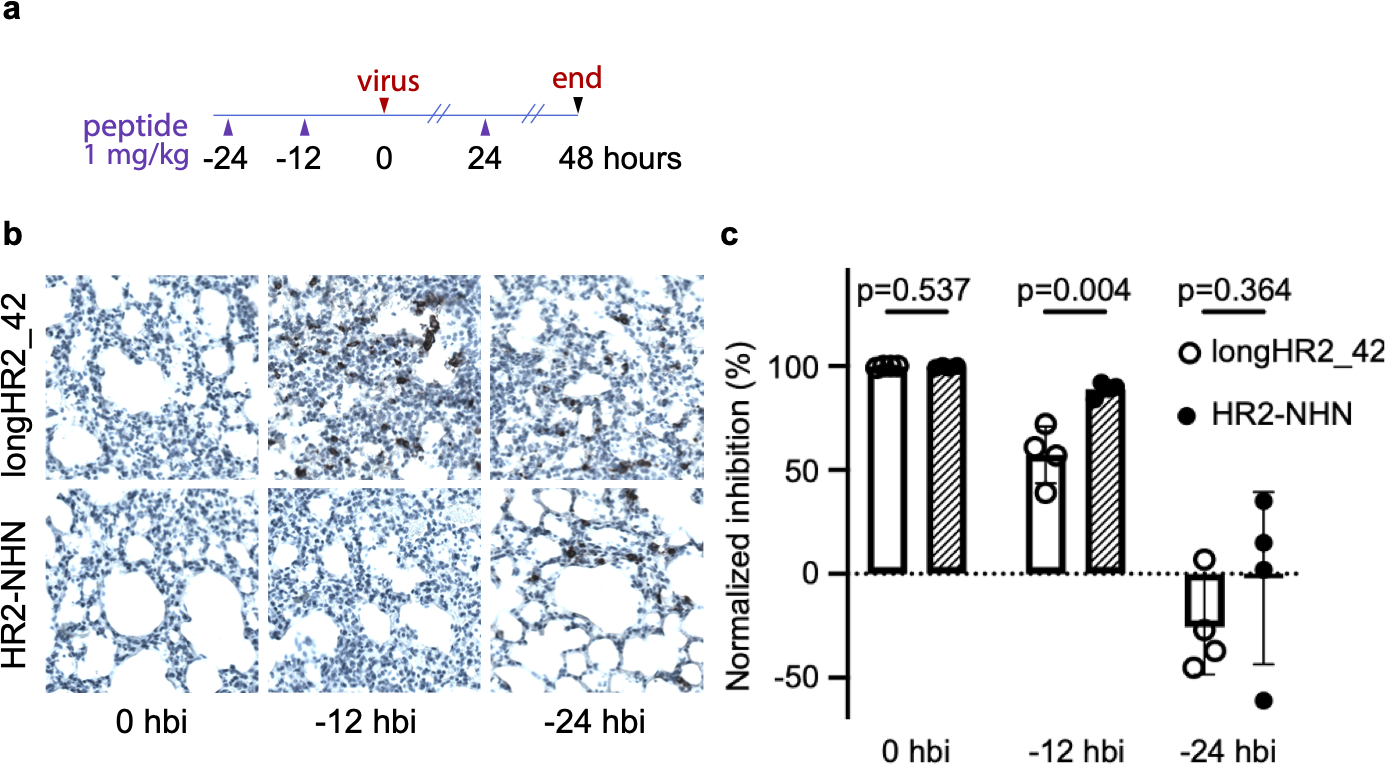

**Fig. S9. The TMPRSS2-resistant HR2-NHN maintains inhibitory activity longer than wildtype longHR2_42 in an authentic SARS-CoV-2 mouse infection model.**

(**a**) Workflow of *in vivo* experiment identifying the duration of antiviral activity for the peptides. (**b**) Example lung section images after immunohistochemistry to visualize viral N protein expression (dark precipitate) accumulating in bronchiolar epithelial cells and pneumocytes counterstained with hematoxylin (light blue staining) at 48 hpi, and (**c**) normalized inhibition quantified by qRT-PCR from the infection of mice with SARS-CoV-2 Beta. In (**b**) and (**c**), infected mice were pretreated with the respective peptides for 0 h, 12 h, and 24 h before infection. In panel (**c**), P values calculated by the unpaired two-tailed t-test are indicated for each time point.

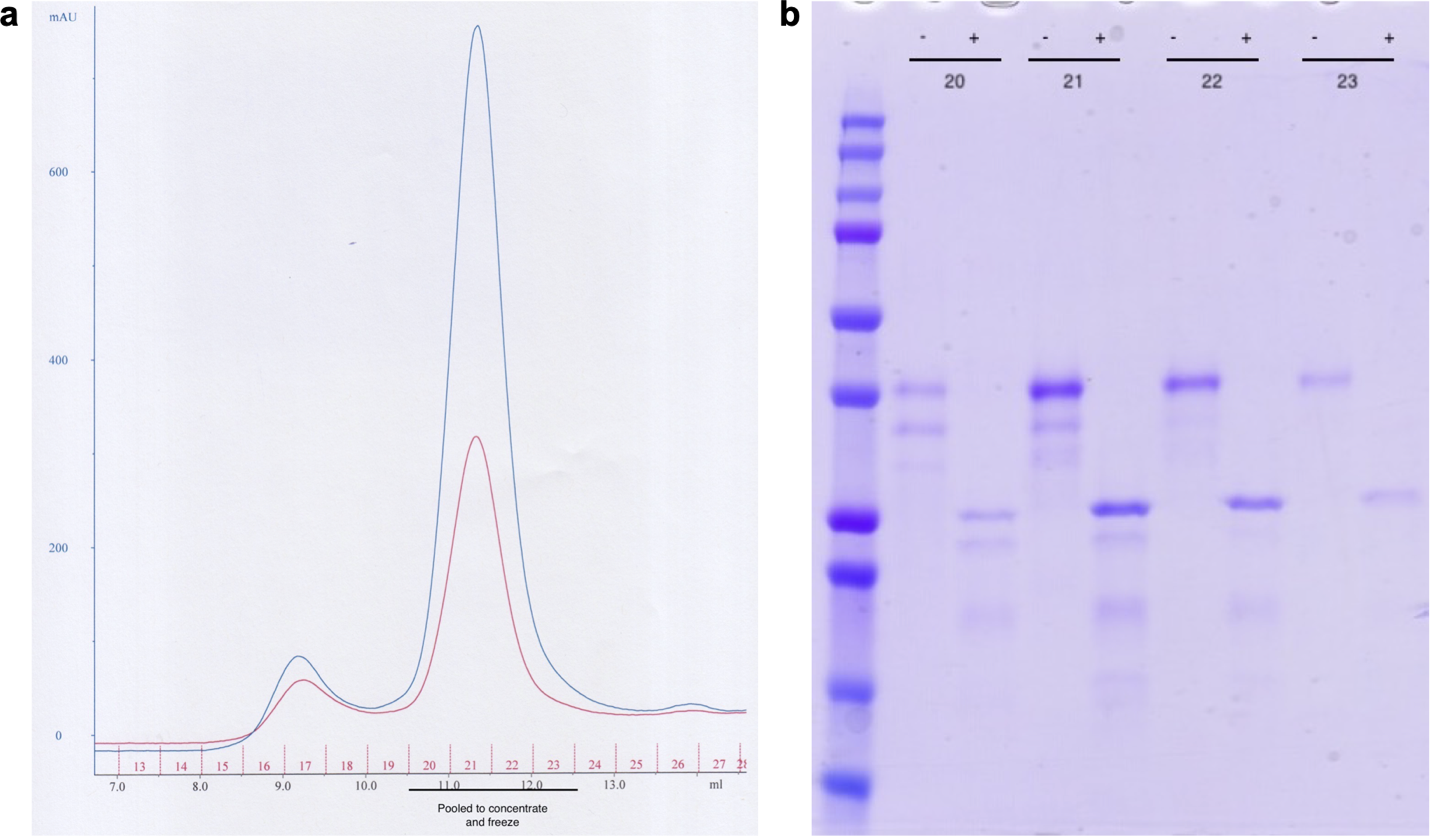

**Fig. S10. Purification of TMPRSS2.**

(**a**) Profile of the last SEC step of a TMPRSS2 preparation. UV absorption curves at 280 nm and 260 nm are colored blue and red, respectively.

(**b**) SDS-PAGE of the pooled fractions in the last SEC step. The fraction numbers were labeled on top of the lanes with the plus and minus signs denoting with and without DTT, respectively.

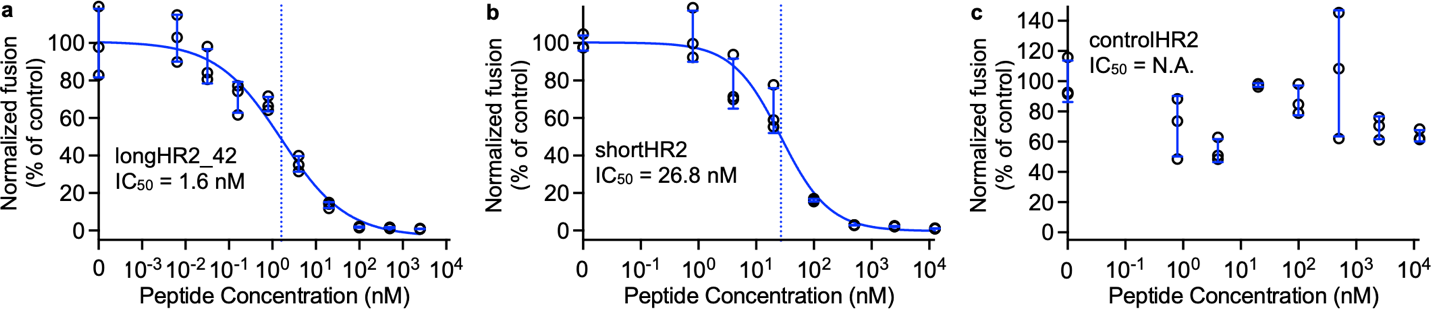

**Fig. S11. Inhibitory activities of peptides prepared with our new protocol in a cell-cell fusion assay.**

The raw data points are plotted as black circles, while the error bars (SD), fitted curves, and vertical dashed lines at IC_50_ are plotted in blue.

**Table S1. Cryo-EM data collection, refinement, and model building**

|  | HR1/HR2-NHN |
| --- | --- |
| EMDB | EMD-70569 |
| PDB | 9okn |
| Microscope | Titan Krios |
| Voltage (kV) | 300 |
| Camera | Gatan K3 |
| Pixel size (Å) | 0.325 |
| Exposure time (s) | 1.1 |
| Number of frames per exposure | 40 |
| Total Dose (e^-^/Å^2^) | 52 |
| Number of movies | 20,031 |
| Defocus range (µm) | -2 to -0.3 |
| Number of particles | 838,766 |
| Resolution of final global refinement (0.143 FSC, Å) | 1.7 |
| Resolution of final local refinement (0.143 FSC, Å) | 2.2 |
| Bond RMSD (Å) | 0.005 |
| Angle RMSD (°) | 0.473 |
| Molprobity score | 1.31 |
| Clashscore, all atoms | 5.72 |
| Ramachandran favored (%) | 98.47 |
| Ramachandran allowed (%) | 1.53 |
| Ramachandran outliers (%) | 0 |
| Rotamer outliers (%) | 0.67 |
| Cβ outliers (%) | 0 |
| CaBLAM outliers (%) | 0 |
